# Modeling the structure-conditioned sequence landscape for large-scale protein design with TriFlow

**DOI:** 10.64898/2025.11.30.691458

**Authors:** Harish Srinivasan, Rongqing Yuan, Jing Zhang, Qian Cong, Jian Zhou

## Abstract

Generative models have revolutionized computational protein design, and the design of high-quality sequences given backbone structure is a critical component for success. Current state-of-the-art design pipelines utilize sequence design methods with local structural context and autoregressive generation. To improve efficiency and quality of sequence design, we developed TriFlow, a model that combines a RoseTTAFold-like three-track architecture for global structural context with discrete flow-matching for efficient few-step sequence generation. We trained TriFlow on a large dataset of interacting protein chains from Protein Data Bank and interacting domains from AlphaFold protein structure Database to enrich its knowledge of natural protein and domain interfaces. TriFlow outperforms existing sequence design methods like ProteinMPNN across diverse benchmarks, including *de novo* binder design, where it boosts the *in silico* success rate of state-of-the-art design pipelines such as BindCraft. We demonstrated this by conducting a large-scale benchmark, generating and computationally validating binders for over 500 diverse protein targets. Experimental validation on a small set of targets also suggests that the performance of TriFlow is on par with BindCraft. By leveraging the model to explore the designed sequence landscape, we discovered that we can effectively highlight functional sites, by contrasting designed sequences that reflect structure constraints with natural evolutionary profiles. As a practical demonstration of its capabilities, we applied our pipeline to systematically design specific binders against human class I cytokines, computationally optimizing for on-target affinity while minimizing off-target interactions, demonstrating that specificity also scales with inference time and computational budget. TriFlow thus provides a robust framework both for large-scale protein engineering and for exploring the fundamental principles of the structure-conditioned sequence landscape. TriFlow software and generated resources are available at https://triflow.zhoulab.io/.

## Introduction

Designing protein sequences that fold into desired three-dimensional structures is a central challenge in computational protein engineering. Modern deep learning based design pipelines typically consist of two stages: backbone generation and sequence design. In the first stage, backbone structures are designed using generative models such as RFdiffusion or hallucination-based approaches like ColabDesign^1,2^. In the second stage, structure-conditioned sequence design models assign amino acids compatible with the target backbone. Finally, structure-prediction tools such as AlphaFold are used to refold the designed sequences and retain only those whose predicted structures closely match the intended backbone^3^.

Recent advances have produced a growing family of sequence design methods. They can be categorized by their core architectures as message-passing neural networks (MPNNs) such as ProteinMPNN and PiFold, transformer-based models such as ESM-IF, and physics-based methods such as Rosetta^4–6^. Among these, MPNNs have historically achieved strong performance when deployed to protein design pipelines, yet their local receptive fields limit their ability to capture long-range structural dependencies^7^.

To overcome these limitations, we developed TriFlow to effectively capture global dependencies across the entire protein or a protein complex by combining advances in generative AI based on discrete flow matching and a new three-track global attention architecture inspired by AlphaFold2 and RoseTTAFold2^3,8^. With discrete flow matching, TriFlow models the global structural context and decodes multiple residues simultaneously in each step, enabling the design of the entire protein sequence in as few as ten inference steps, regardless of protein size^9^. Trained on Protein Data Bank (PDB) structures using the same data splits as ProteinMPNN, TriFlow achieves superior performance across both monomer and binder design benchmarks. Moreover, we scaled the model training by creating a larger dataset made of interacting domain pairs in AlphaFold models, to improve design accuracy and generalization^10^.

Leveraging the efficiency and robust performance of TriFlow, we conducted several large-scale *in silico* experiments to gain insights into the quality, specificity, and properties of designed sequences. We performed a large-scale *in silico* binder design campaign across more than 500 randomly selected protein targets. TriFlow consistently outperformed ProteinMPNN as a drop-in replacement in state-of-the-art design pipelines. Our analysis also uncovered factors influencing binder design success, including the chemical properties and secondary structures of interface residues, as well as the size of the endogenous interface.

A critical challenge in deploying designed binders for biomedical applications is minimizing off-target effects. We reason that an efficient sequence designer such as TriFlow can enable effective exploration of sequence and structure space, thereby facilitating the identification of binders with enhanced specificity. Using human class I cytokines as a model system, we performed an *in silico* evaluation of binder specificity and sought to generate selective binders for each member of the human class I cytokine family.

Although off-target interactions were prevalent, we found that they can be substantially reduced by *in silico* selections of designed sequences against undesired binding. The designed specific binder sequences for human class I cytokines are released as a public resource to support potential biomedical applications.

Finally, we designed sequences for a set of 44k diverse domains representing all known homologous folds in Evolutionary Classification Of protein Domains (ECOD)^11^. Comparing the designed and natural sequence profiles showed the largest disparities at protein active sites and small-molecule binding pockets. These findings point to a new use of structure-conditioned sequence design for large-scale functional annotation and suggest that jointly designing structure and function will be an important future challenge in protein engineering.

## Results

### TriFlow: modeling global context with discrete flow and global attention

To capture the full context of the protein structure for the sequence design problem, we combined discrete flow matching with a three-track evoformer-like architecture. Unlike autoregressive models such as MPNNs, discrete flow matching captures global dependencies and enables parallel decoding of multiple amino acid positions, allowing few-step sequence design for any backbone size (**Fig. 1a**). The three-track architecture utilizes transformers to integrate the single, pair, and 3D representations and capture both short and long-range interactions in proteins (**Fig. 1b**). Unlike the RoseTTAFold architecture that exploits biased axial attention for the pairwise updates, we use invariant point attention and only use triangle multiplicative updates (Supplementary **Fig. S1**) to allow direct reasoning about 3D geometry and improve computational efficiency.

**Figure 1.**
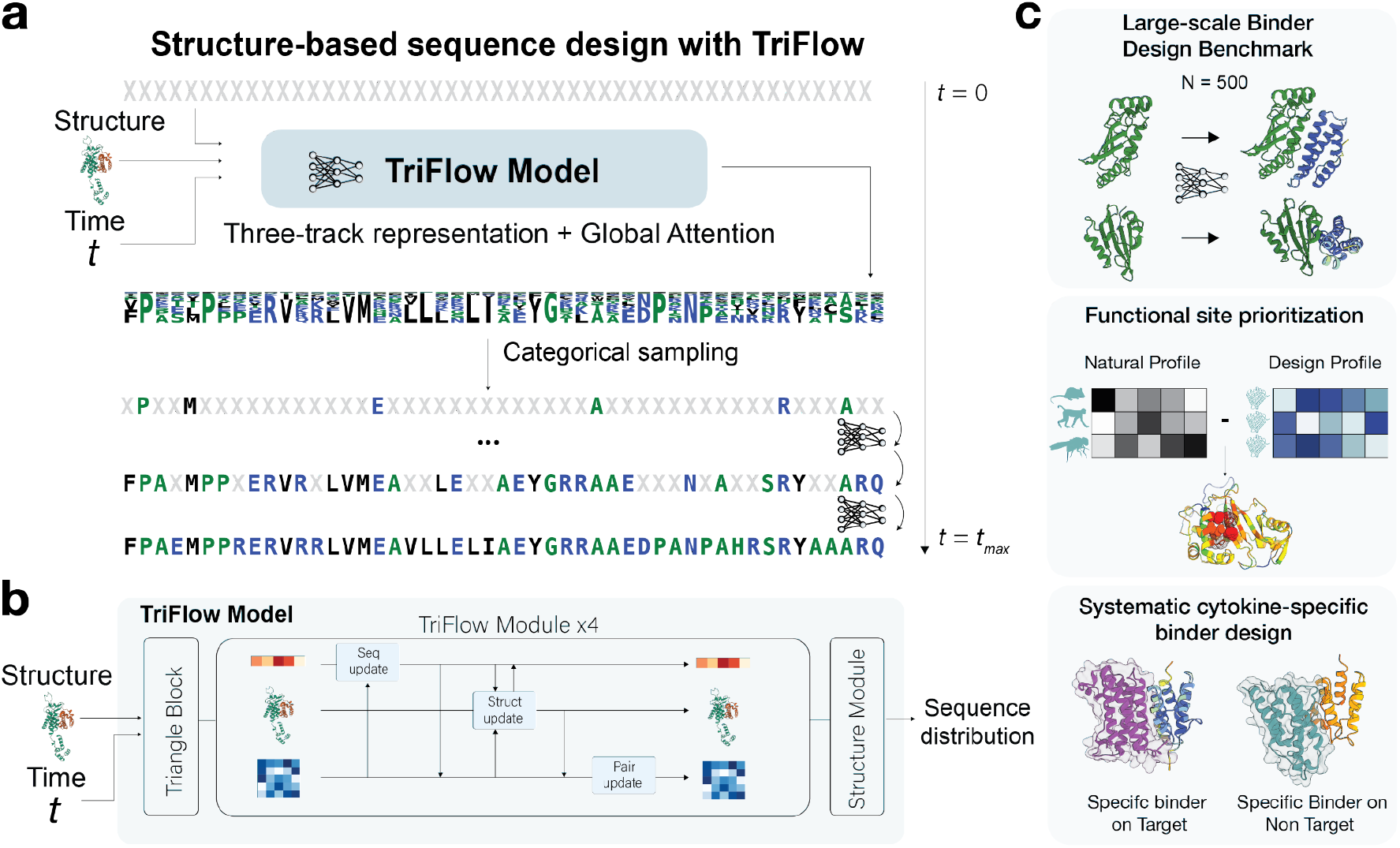
TriFlow overview. **a)** TriFlow sequence generation process starts with all masked tokens, and the model predicts a probability distribution for each amino acid position. These are sampled at each step to unmask the sequence until t_max_. **b)** Key components in the TriFlow model (more details in Supplementary **Figure S1**), which takes backbone structure, timestep, and sequence from a previous time step as input. **c)** Applications of our sequence design model in large-scale binder design benchmark, functional site prediction, and specific binder design for human cytokines.

As we anticipated that model performance would scale with training data size, we trained TriFlow on two datasets: a PDB dataset with the same train–test splits as ProteinMPNN to enable direct comparison, and an extended distilled datasets derived from predicted structures in the AlphaFold protein structure Database (AFDB). In addition to PDB and AFDB monomer structures, we further included interacting PDB chains as well as interacting domain–domain pairs predicted by AFDB^12^ to strengthen TriFlow’s ability to design protein binders. To our knowledge, ESM-IF is the only structure-conditioned sequence design model trained on AFDB monomers, and no existing models have been trained on a dataset of predicted domain–domain interactions.

However, direct training on the expanded dataset that included AlphaFold-predicted structures did not improve TriFlow’s sequence generation performance on experimental backbones. We hypothesized that inaccuracies in AlphaFold-generated 3D coordinates, particularly in regions with lower prediction confidence, may limit their utility for training structure-conditioned sequence design models. Previous studies have shown that introducing structural noise during training can improve robustness and performance in protein design applications^4^. Building on this idea, we trained the model with multiple noise levels, enabling it to account for varying degrees of local uncertainty in backbone coordinates. Incorporating noise proved critical for effectively leveraging AlphaFold models during training. With this strategy, the model trained on the expanded dataset achieved performance comparable to that of the model trained exclusively on experimental structures (Supplementary **Fig. S2**).

We extensively benchmarked TriFlow performance, including creating a new large-scale binder-design benchmark with 500 diverse target proteins, and demonstrated several novel applications of sequence design, including prediction of functional sites and designing specific binders for human class I cytokines (**Fig. 1c**, and details below). To compare performance between TriFlow and other methods, we used the TriFlow version trained on the ProteinMPNN training set, and for the large-scale binder design and other applications, we used the TriFlow version trained on the extended dataset of both PDB structures and AFDB models.

### TriFlow improves protein sequence design

We performed comprehensive evaluations of TriFlow’s ability in sequence recovery and protein monomer design. First, we evaluated TriFlow’s ability to recover sequences for given structures using the shared PDB test set of 1392 structures between ProteinMPNN and TriFlow. TriFlow shows higher sequence recovery rate than previous methods, including ProteinMPNN, ESM-IF, and PiFold^4–6^ (**Fig. 2a**). Second, to test TriFlow’s ability to design sequences for generated backbones, we used 3D structures with sequence lengths ranging from 100 to 1000, generated by a hallucination method using the ColabDesign framework built on top of AlphaFold^13^. For each backbone size, we evaluated the refoldability of designed sequences, as indicated by the highest TM score between the target backbone and the predicted 3D structure of the designed sequence by AlphaFold3 in a single-sequence mode^14,15^. TriFlow outperforms all other models in sequence refoldability, particularly at the longer lengths, ranging from 600 to 1000 residues (**Fig. 2b**).

**Figure 2.**
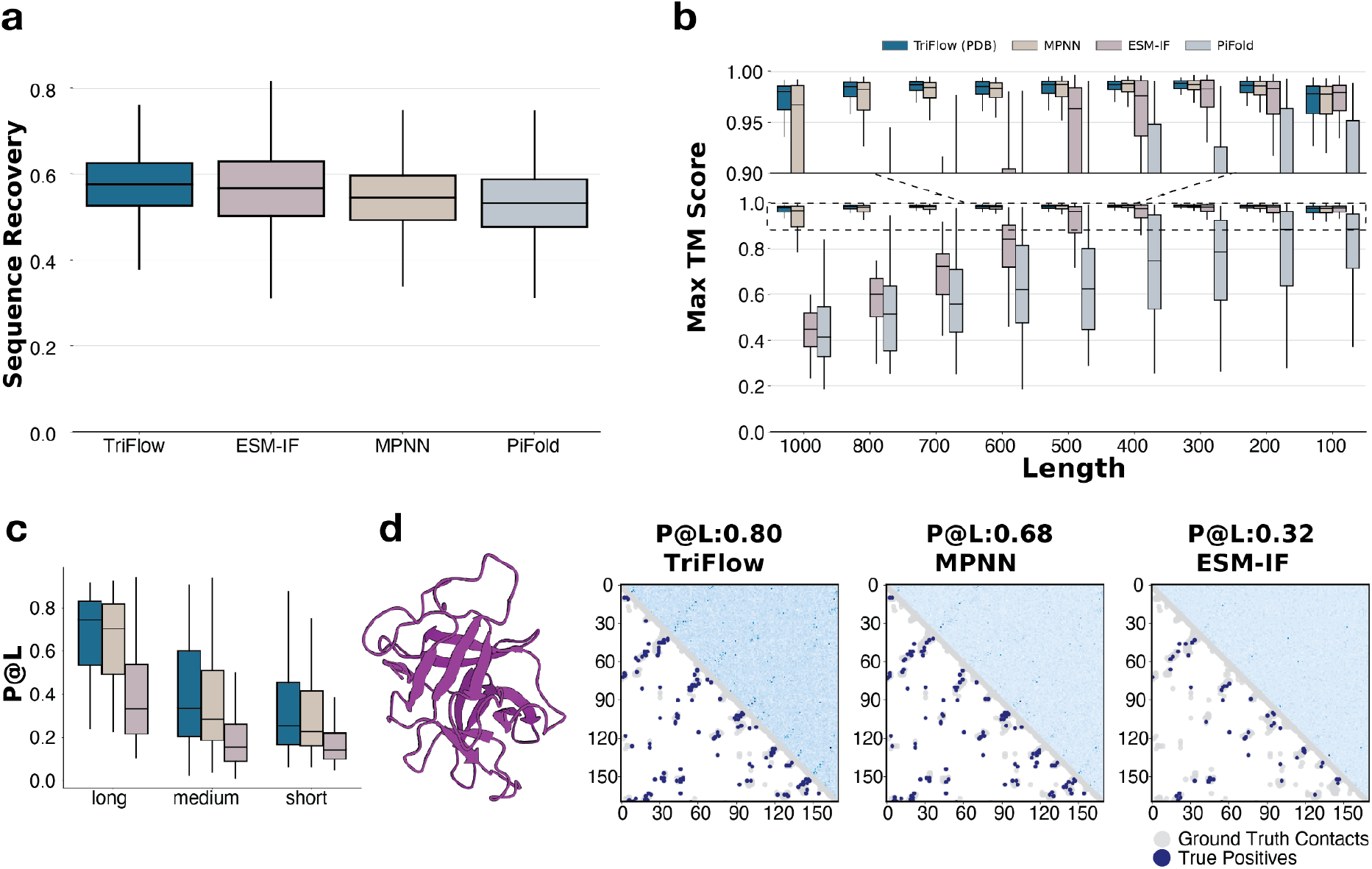
TriFlow outperforms previous tools in monomer sequence design. **a)** Native sequence recovery across different tools evaluated on the common test set used by ProteinMPNN and TriFlow, calculated as the fraction of residues for which the designed and natural sequences share the same amino acid. Potential train–test overlap cannot be excluded for ESM-IF and PiFold, which may slightly inflate their reported performance. **b)** Refoldability of sequences generated by different methods for a collection of ColabDesign-generated backbones of varying sizes (100–1000 residues)13. Refoldability is quantified as the maximal TM-score between the original backbone and the structures predicted by AlphaFold3 for the designed sequences. **c)** Agreement between the top-L coevolving residue pairs (L = protein length), inferred using GREMLIN16, and the inter-residue contacts in the native 3D structures. The analysis was performed on 30 representative proteins (clustered at 30% sequence identity) from the ProteinMPNN/TriFlow test set. Residue pairs were stratified by sequence separation: short (6 ≤ separation < 12), medium (12 ≤ separation < 24), and long (separation ≥ 24). **d)** Example protein structure (left) and the agreement between native contacts and the top-L coevolving residue pairs inferred from sequences designed by different tools. For each method, the upper triangle shows the coevolutionary couplings between residues inferred from the generated sequences, and the lower triangle shows the ground-truth contacts in the native structure. True positives that appear both among the top-L coevolving pairs and in the ground-truth contacts are highlighted in blue, and the remaining native contacts shown in grey.

Interestingly, we found that introducing noise into backbone structures decreases TriFlow’s sequence recovery rate but improves the refoldability of the designed sequences, particularly for larger proteins (Supplementary **Fig. S3**). A similar tradeoff has been reported for previous sequence design models, including ProteinMPNN. We hypothesize that noise augmentation reduces the model’s reliance on fine-grained local backbone geometries that are informative for recovering the native sequence, thereby lowering sequence recovery. At the same time, it encourages the model to learn sequence–structure relationships that are robust to local structural inaccuracies, preventing overfitting to backbone errors. As a result, the designed sequences exhibit improved refoldability, especially for longer proteins, where local backbone inaccuracies are more prevalent and can have a larger cumulative impact on the global fold.

We reasoned that TriFlow’s use of global structural context may enable it to more effectively capture dependencies between distant positions. Such inter-residue dependencies are typically reflected in amino acid covariance or coevolution signals, which in natural sequences arise from pairwise evolutionary constraints and closely mirror physical contacts within protein structures^17^. To evaluate whether designed sequences conditioned by structures satisfy similar coevolutionary constraints, we randomly selected 30 proteins from distinct clusters in the test set and generated 1,000 sequences for each. Using GREMLIN, we quantified residue–residue “coevolution” and assessed whether residue pairs with the strongest coevolutionary signals (top L, L/2, and L/5 pairs) correspond to inter-residue contacts in the 3D structures¹³ (**Fig. 2c-d**).

We compared TriFlow with ProteinMPNN and ESM-IF; PiFold was excluded because it deterministically produces only a single sequence per backbone and is not suitable for this task. We stratified inter-residue contacts by sequence separation into short, medium, and long-range categories. Across all categories, co-varying/evolving residues in TriFlow-designed sequences show the highest agreement with structural contacts (**Fig. 2c**). This suggests that TriFlow captures amino acid dependencies more effectively than previous models, indicating a stronger ability to learn structural constraints that govern residue–residue interactions. Inspection of a representative example reveals that the sequence profiles generated by ESM-IF show little correspondence with residue contacts in the native 3D structure (**Fig. 2d**). In contrast, both TriFlow and ProteinMPNN recover coevolutionary signals associated with structural contacts. Notably, TriFlow exhibits a stronger correspondence between its sequence profiles and the underlying structural contact map, suggesting a more accurate representation of structure-derived residue dependencies.

Another factor that could influence the coevolutionary signals in the generated sequences is the sequence diversity. This diversity is controlled by the *temperature* parameter, which determines the sharpness of the sampling distribution. The analyses above use the default temperature of 0.1 across all methods. We also ablated the temperature values from 0.1 to 1 to generate sequences of varying diversity and to observe their effects on amino acid dependencies between positions (Supplementary **Fig. S4**). All three models improved as temperature increased and achieved their best performance at a temperature near 1.0. Notably, TriFlow recovered structure-associated inter-residue dependencies more effectively than the other methods even at lower temperatures. This superior ability to capture structural constraints may also contribute to its superior performance in sequence recovery and refoldability.

### TriFlow improves state-of-the-art *de novo* binder design pipelines

Targeting natural proteins with designed binders is a major application of *de novo* protein design. Binder-design pipelines typically follow four stages (**Fig. 3a**): selecting a hotspot or interface on the target (optional), generating a binder backbone conditioned on that site, designing a sequence compatible with the target–binder complex, and finally refolding the complex using a structure-prediction model such as AlphaFold3 in single-sequence mode with the target chain supplied as a template (**Methods)**^18,19^. Across nearly all contemporary pipelines, ProteinMPNN has been used as the standard model for sequence design^1,2,13,20–22^. Two widely used approaches for backbone design are RFdiffusion and ColabDesign; the former designs binders using hotspots sampled from native interface, whereas the latter does not require pre-defined hotspots to guide the structure generation process^1,2^.

**Figure 3.**
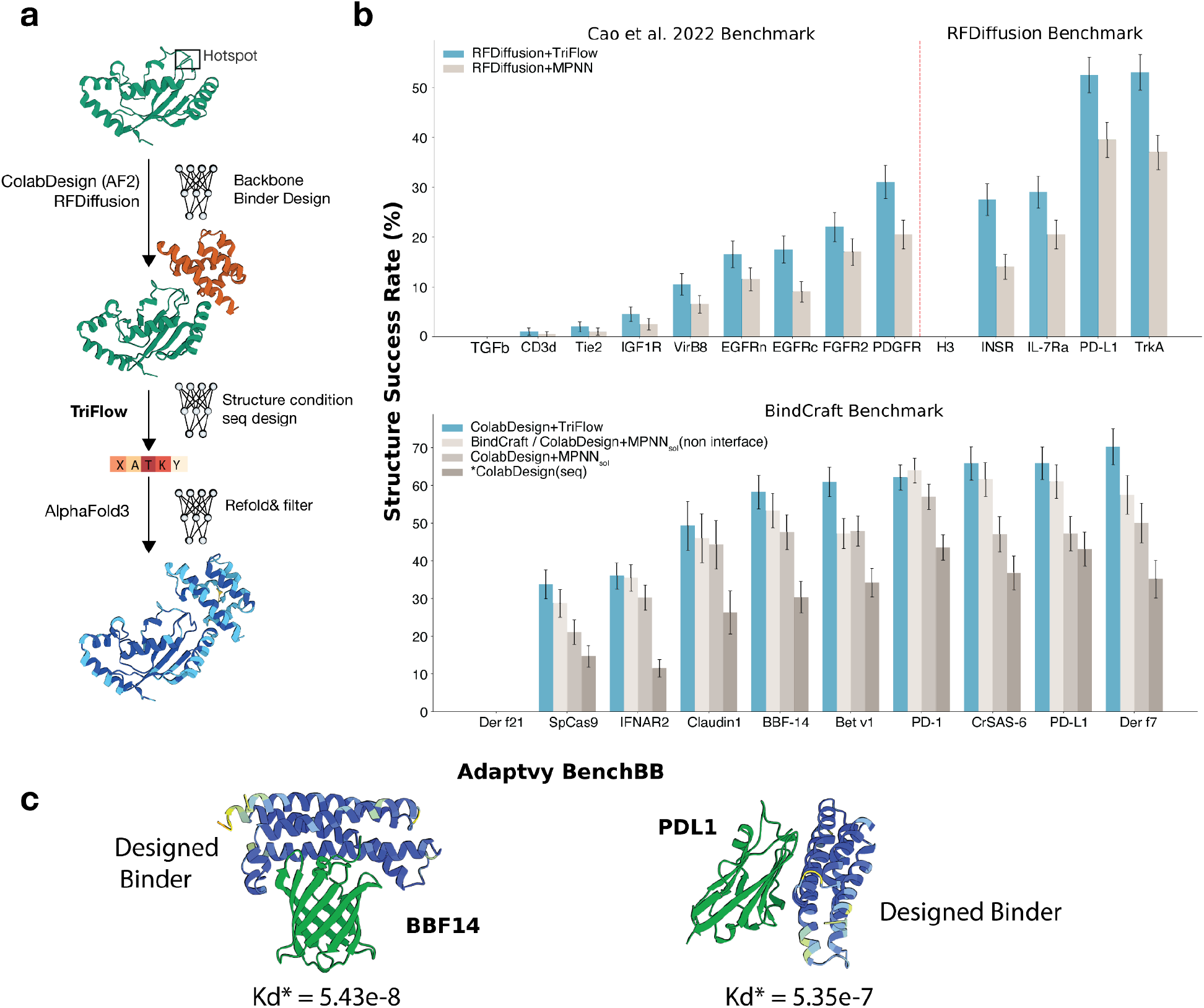
TriFlow outperforms ProteinMPNN and SolubleMPNN in protein binder design. **a)** Workflow describing the binder design pipeline: we first design backbones with RFdiffusion and ColabDesign, respectively, with binding hotspots specified for RFdiffusion; second, a set of sequences is generated with TriFlow for each binder backbone and then refolded with AlphaFold3. **b)** Performance of TriFlow and MPNN-based models evaluated by structure success rate, i.e., the percentage of binder backbones with at least one designed sequence that passed the established cutoffs based on AlphaFold3 confidence metrics. *Top:* Structure success rate on the combined set of therapeutic targets from a previous *de novo* binder-design study and the RFdiffusion paper. RFdiffusion+TriFlow: backbones generated by RFdiffusion with 8 sequences designed by TriFlow per backbone. RFdiffusion+MPNN: backbones generated by RFdiffusion with 8 sequences designed by ProteinMPNN per backbone. *Bottom:* Structure success rate on the BindCraft benchmark set. ColabDesign+TriFlow: backbones generated by ColabDesign with 8 sequences designed by TriFlow per backbone. BindCraft: backbones generated by ColabDesign with 8 sequences designed by SolubleMPNN per backbone, retaining the ColabDesign interface residues following the original BindCraft protocol. ColabDesign+MPNN_sol_: backbones generated by ColabDesign with 8 sequences designed by SolubleMPNN, including redesign of interface residues. *ColabDesign: both backbone and sequence generated by ColabDesign, which produces a single deterministic sequence per backbone and frequently performs poorly in experimental validation due to poor expression. **c)** Predicted structures of Triflow-designed binders bound to their respective targets, BBF14 (left) and PD-L1 (right). Designed binders are colored blue, and target proteins are colored green.

To evaluate TriFlow in realistic *de novo* binder design settings, we benchmarked it against a panel of 24 therapeutically relevant protein targets from previous studies that used the RFdiffusion and BindCraft pipelines^1,2,23^. We first compared TriFlow’s sequence design ability against the original ProteinMPNN on RFdiffusion generated structures and a modified variant, SolubleMPNN (or MPNN_sol_), on BindCraft generated structures^4,20^.

We quantified performance using two metrics: the **structure success rate**, defined as the percentage of designed binder backbones that passed the established AlphaFold3 confidence thresholds after refolding (see **Methods**), and the **sequence success rate**, defined as the fraction of designed sequences that satisfied the same refolding criteria. We primarily relied on structure success rate because diversifying the scaffolds in the final pool of designed proteins are preferred in downstream experimental tests and sequence optimization^1^.

Across all designed backbones (100-200 each) for 24 targets, we designed 8 sequences with each tool and evaluated them with AlphaFold3. TriFlow achieves a higher structure success rate (**Fig. 3b**) than either ProteinMPNN or SolubleMPNN. Three of the 24 targets (TGFβ, H3, and Der f21) could not be evaluated due to technical limitations in handling multi-chain templates in the AlphaFold3-based binder-evaluation script. Likewise, TriFlow outperforms both ProteinMPNN and SolubleMPNN in sequence success rate (see **Methods**) for 19 of the 21 targets (Supplementary **Fig. S5** and **S6**). The only two exceptions are Claudin1 and PD-1 (Supplementary **Fig. S6**), for which all three methods yield comparable performance. Together, these results demonstrate that TriFlow consistently generates higher-quality binder sequences than ProteinMPNN.

The widely used BindCraft pipeline combines ColabDesign with SolubleMPNN and it represents the state-of-the-art for protein binder design. ColabDesign uses an AlphaFold2-based hallucination framework. Starting from a randomly initialized binder, gradients were propagated through AlphaFold2 predictions to iteratively optimize sequences toward designs predicted to fold into stable structures and form favorable interfaces with the target protein. Sequences designed by ColabDesign are thus directly optimized for AlphaFold confidence metrics^1,24^. However, although ColabDesign-generated sequences show high AlphaFold3 confidence (Supplementary **Fig. S6**), they often suffer from poor expression and solubility in experimental settings because the optimization is only based on structure loss and does not take the necessary sequence distribution, such as solvent accessibility surface and charge distribution, into consideration. Subsequent redesign with SolubleMPNN has been shown to substantially improve experimental success rates^1,20^. Preserving interface residues designed by ColabDesign while redesigning only the non-interface residues with SolubleMPNN further boosts the structure success rate.

Replacing SolubleMPNN with TriFlow improved the structure success rate for all but one target (PD-1) in the BindCraft benchmark (**Fig. 3b, bottom**). Notably, this improvement does not depend on retaining the interface residues originally designed by ColabDesign and optimized for AlphaFold2 confidence metrics. In fact, redesigning **all** residues including interface positions with TriFlow yielded the highest structure success rate (Supplementary **Fig. S6**), as measured by AlphaFold3 refoldability. This result indicates that TriFlow’s interface design is as effective as direct optimization through AlphaFold2.

To experimentally evaluate the performance of TriFlow, we tested designed binders against three targets—PD-L1, BBF14, and Cas9—using Bio-Layer Interferometry (BLI) and Surface Plasmon Resonance (SPR) through Adaptyv’s BenchBB^25^ framework. For each target, we selected the top-ranked designs based on ipAE_min_ from five independent backbones generated with ColabDesign. As a baseline, we similarly selected the best designs from five backbones generated using the BindCraft pipeline, which combines ColabDesign backbone generation with SolubleMPNN redesign of non-interface residues. The ColabDesign–TriFlow pipeline yielded three experimentally validated binders across two of the three targets tested, including two binders against PD-L1 and one against BBF14 (**Fig. S7**), displaying the same experimental success rate as the established BindCraft pipeline. BLI and SPR sensorgrams confirmed robust binding of the validated TriFlow designs to their respective targets, with affinities comparable to those of the corresponding BindCraft (SolubleMPNN) designs (**Fig. S8**). The strongest TriFlow binders, shown in **Fig. 3c**, exhibited dissociation constants (Kd) of 5.35 × 10⁻⁷ M for PD-L1 and 5.43 × 10⁻⁸ M for BBF14, corresponding to submicromolar and nanomolar binding affinities, respectively. Overall, TriFlow achieved experimental performance comparable to that of ProteinMPNN-based design while producing high-affinity binders against two of the three target proteins tested, despite screening only a small number of candidate designs experimentally.

### Large-scale binder design benchmark

To comprehensively evaluate model performance across a broad spectrum of protein targets and understand factors that are linked to successful binder design, we constructed a large-scale binder-design (LBD) benchmark consisting of 500 diverse binary complexes randomly sampled from PDB entries with low (< 30%) sequence identity to proteins in the shared TriFlow–ProteinMPNN training dataset. This benchmark was designed to evaluate how TriFlow performs, instead of MPNN-based networks, within two widely adopted binder-design pipelines: (1) RFdiffusion → ProteinMPNN and (2) ColabDesign → SolubleMPNN (i.e., the BindCraft pipeline).

For each target, we generated 25 binder backbones using RFdiffusion and up to 10 backbones using ColabDesign adapted from the BindCraft pipeline (see **Methods**). For each RFdiffusion-generated backbone, we designed 8 sequences using TriFlow and 8 using ProteinMPNN. For each ColabDesign-generated backbone, we designed 8 sequences using TriFlow and 8 using SolubleMPNN; TriFlow redesigned all residues, whereas SolubleMPNN redesigned only non-interface positions, following the practice of the BindCraft pipeline. In total, this benchmark comprises 16,216 binder backbones (12,500 from RFdiffusion and 3,716 from ColabDesign) and 129,728 designed sequences.

Across the entire LBD benchmark, TriFlow consistently outperforms ProteinMPNN and SolubleMPNN. For RFdiffusion-generated backbones, TriFlow achieves a higher structure success rate in 143 targets, whereas ProteinMPNN performs better on only 68 (**Fig. 4a** left). For ColabDesign-generated backbones, TriFlow performs better on 200 targets, while SolubleMPNN outperforms TriFlow on only 93 (**Fig. 4a** right). It is important to note that we designed only 8 sequences per backbone for the LBD benchmark, meaning the measured success rates for individual targets are subject to stochastic variability. We expect TriFlow’s advantage to become even more pronounced when more sequences per backbone are generated, as observed in our earlier small-scale benchmarks (**Fig. 3b**).

**Figure 4.**
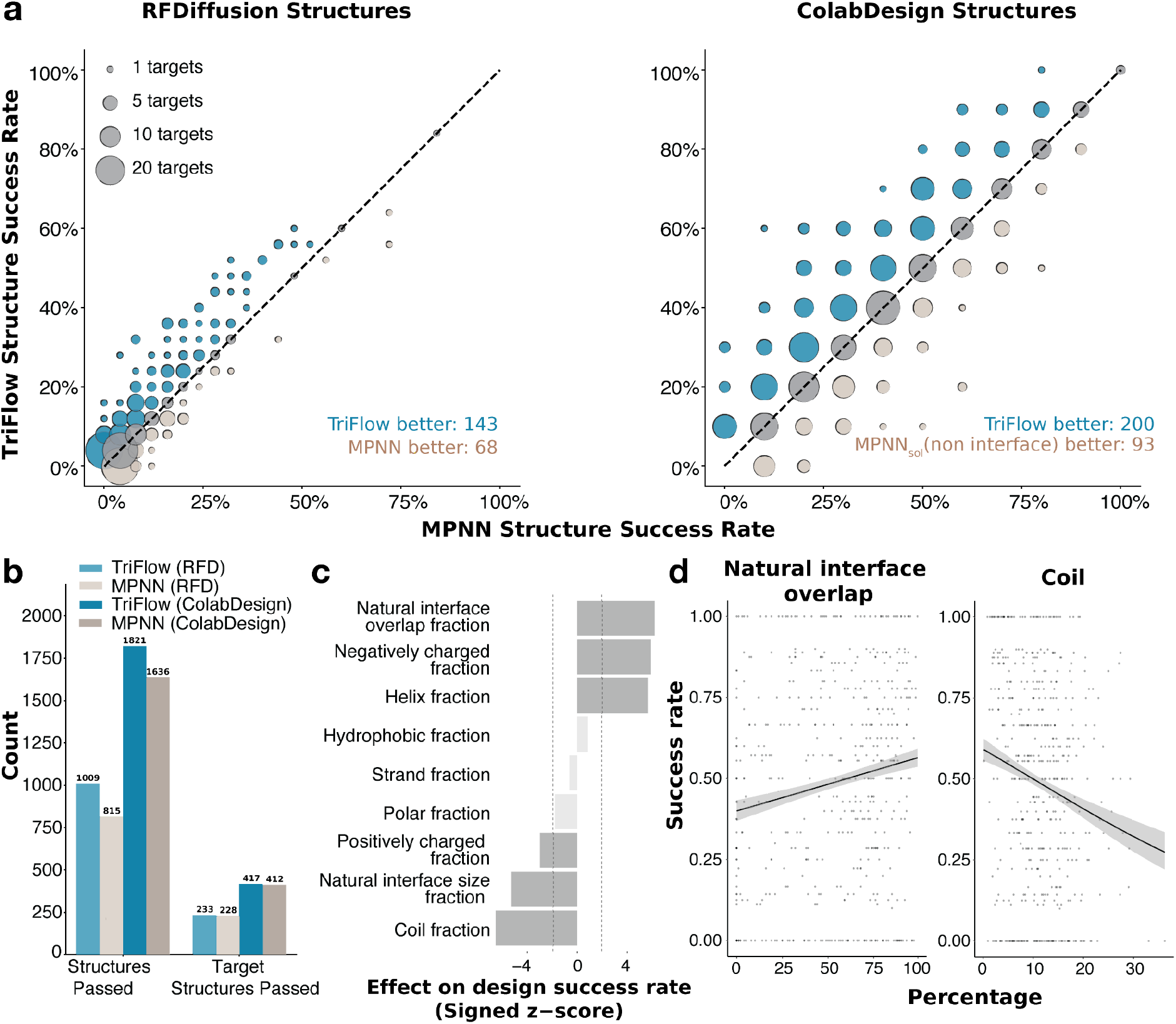
Large-scale benchmark of binder design pipelines. **a)** Comparison of structure success rates by different pipelines on 500 target structures. Left: binder backbones generated by RFdiffusion and 8 sequences designed by TriFlow and ProteinMPNN, respectively, for each backbone. Right: binder backbones generated by ColabDesign and 8 sequences designed by TriFlow and SolubleMPNN, respectively, for each backbone. **b)** Summary of the total number of binders that pass the *in silico* evaluation criteria (see **Methods**) for each pipeline, along with the number of targets for which at least one successful binder is obtained. In the labels above, the structure design method is indicated in parentheses and the sequence design method is outside the parentheses; RFD denotes RFdiffusion. **c)** Z-scores of logistic-regression coefficients quantifying the association between target features and design success. Positive values indicate that higher feature values are associated with increased success rates, and negative values indicate that higher feature feature values decrease success rate. **d)** Scatterplots with logistic regression fits showing the relationship between structure success rate and two target properties: (i) interface overlap between the designed and natural binders, and (ii) the proportion of coil residues at the target interface engaged by the designed binders.

Both TriFlow and MPNN perform better on ColabDesign backbones, yielding successful designs for 1,821 | 1,636 (49% | 44%) ColabDesign backbones across 417 | 412 (83% | 82%) targets, compared with 1,009 | 815 (8.0% | 6.5%) RFdiffusion backbones across 233 | 228 (47% | 46%) targets. (**Fig. 4b**). We compared structural properties of successfully designed binders generated by RFdiffusion versus ColabDesign against native binders of our targets. Since RFdiffusion is guided by the native hotspots to reuse the native interface, ColabDesign was not guided to use the native interface, but binders designed by it still show large overlap with native interface residues (Supplementary **Fig. S9a**).

The LBD benchmark consists of randomly selected binary protein complexes from the PDB. Compared with interfaces in multi-component protein assemblies, these complexes are enriched for transient or non-obligate interactions, which typically have smaller buried interface areas than obligate interactions within stable multi-protein assemblies^26^. Binders designed by both RFdiffusion and ColabDesign form larger interfaces with their targets than the corresponding native binders (Supplementary **Fig. S9b,c**). In addition, both structure-generation methods predominantly produce α-helical scaffolds with minimal β-sheet content. However, binders generated by ColabDesign contain a higher proportion of coil regions than those generated by RFdiffusion (Supplementary **Fig. S10a**). Consistent with the scarcity of β-sheet-containing designs, both methods achieve higher success rates on α-helical targets than on β-strand-rich targets, with this structural bias being more pronounced for RFdiffusion-generated backbones (Supplementary **Fig. S10b**).

To better understand which target properties influence sequence-design success, we leveraged our LBD benchmark to correlate diverse target features with design performance using the ColabDesign→TriFlow pipeline. We found that information about the native binding interfaces on a target are useful in guiding protein design strategies. These interfaces, shaped by evolutionary pressure, tend to have the physicochemical and geometric properties that are inherently suitable for binding^27^. Therefore, we observed that greater overlap between the designed binder interface and the native interface on the target correlates with higher success rates (**Fig. 4c** and **4d** left). This observation supports the widely used strategy of anchoring *de novo* binders at native binding sites. Additionally, we found that the native interface size negatively correlates with design success (**Fig. 4c**). This effect is possibly an artifact due to the binder-size constraint imposed during backbone generation (60–120 residues). For targets with large native interfaces, the designed binders are likely too small to occupy the full binding site, making it more difficult for AlphaFold3 to identify a stable, correct binding mode for the designed complex. To improve binder design success rate, it might be important to scale binder size with the native interface size.

In addition, we found that properties of the target interfaces engaged by the designed binders are also predictive of design success. First, target interfaces with higher α-helical content are associated with higher success rates, whereas interfaces enriched in coil regions show lower success (**Fig. 4c**, **Fig. 4d right,** and Supplementary **Fig. S11**). This likely reflects the fact that designed binders are themselves highly α-helical, making them more compatible with helical interface geometries. Second, target interfaces with a larger proportion of negatively charged residues are associated with greater success, whereas interfaces enriched in positively charged residues tend to be more challenging (**Fig. 4c**, and Supplementary **Fig. S11**). We reason that this trend may also arise from the helical nature of the designed binders: positively charged residues (Arg and Lys) have strong helical propensities, whereas one of the negatively charged residues, Asp, is a well-known helix breaker^28,29^. Thus, designing positively charged helical binders for negatively charged interfaces is relatively straightforward, whereas designing negatively charged helical binders for positively charged interfaces is more challenging. Finally, we observed a modest positive correlation between interface hydrophobicity and design success. Because native protein–protein interfaces typically feature hydrophobic residues at the core and hydrophilic residues at the periphery, the relationship between hydrophobicity and successful design is likely more nuanced than a simple monotonic trend^30^.

### Designing specific binders for all class I cytokines

While prior *de novo* design studies have successfully generated binders for individual targets, few have developed systematic pipelines to evaluate and minimize off-target binding to closely related proteins. To explore possible strategies to design highly specific binders, we applied the ColabDesign→TriFlow pipeline to the human class I cytokines, which comprises 31 homologous proteins. Class I cytokines are four-α-helix bundle signaling proteins that act through WSXWS-containing hematopoietin receptors to regulate immunity, development, and hematopoiesis^31^. Class I cytokines are major therapeutic targets, and their dysregulation contributes directly to autoimmune diseases, inflammation, immunodeficiency, cancer, and blood disorders^32^. Existing therapeutics for Class I cytokines are predominantly protein-based agents that bind the cytokines and prevent them from engaging their receptors and initiating downstream signaling^33^. Because Class I cytokines act extracellularly, they are readily accessible to biologics without the challenges associated with intracellular delivery^33^. These properties make Class I cytokines particularly attractive targets for *de novo* designed protein therapeutics.

Using Class I cytokines as an example, we set out to evaluate the target-specificity of *de novo* designed binders. For each cytokine target, we generated up to 10 backbone structures using ColabDesign, designed 40 sequences per backbone with TriFlow, and screened with AlphaFold3 to identify highly confident *de novo* binders. We were able to obtain binders for all cytokine targets except for IL7.

To evaluate the off-target potential of each designed binder, we used AlphaFold3 to model its interactions with every class I cytokine (**Fig. 5a** and Supplementary **Fig. S12a,b**). We quantified binder specificity using a specificity score, defined as the difference between the lowest predicted ipAE_min_ for the intended target (on-target) and the lowest predicted ipAE_min_ among all off-target cytokines (see **Methods**). For most cytokines, we identified multiple binders with high specificity scores (Supplementary **Fig. S12c**). We reasoned that target specificity could be further improved by generating more sequences for each backbone with TriFlow and selecting those with the highest specificity scores, taking advantage of TriFlow’s rapid inference (seconds per sequence versus hours per backbone for ColabDesign). Consistent with this expectation, the best specificity score for each target increased as more sequences were generated (**Fig. 5b**), with the largest improvements observed when increasing the number of sequences from 1 to 30 per target. Beyond 30 sequences, specificity scores largely plateaued for most cytokines, with the exceptions of IL-6 and EPO. Using this simple scaling strategy, we obtained highly specific binders for most cytokines. In contrast, specificity for some targets, particularly GH2, plateaued at moderate levels, suggesting that achieving higher specificity for these proteins may require additional backbone generation or alternative interface hotspot selection.

**Figure 5.**
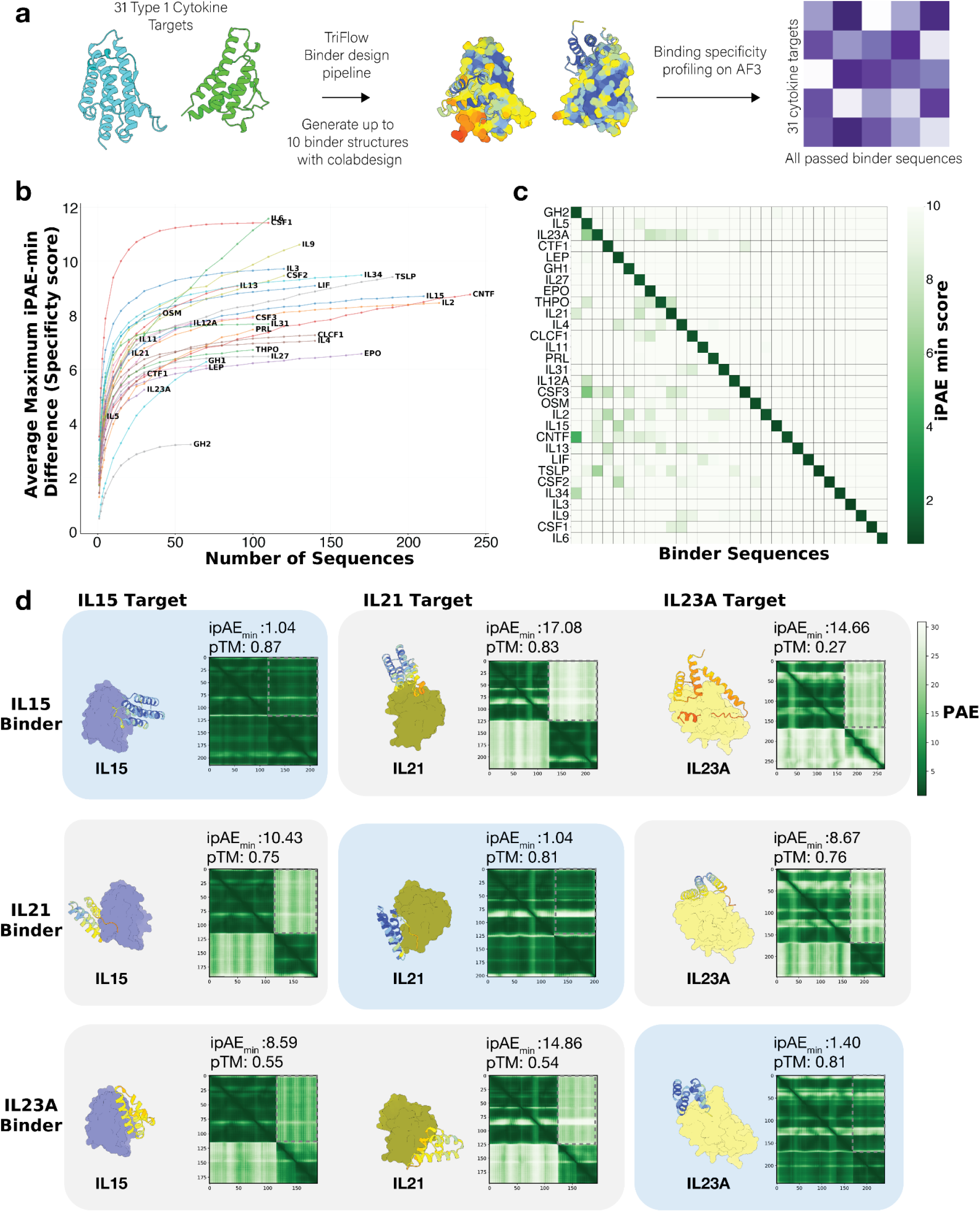
Systematic design of specific binders for class I cytokines. **a)** Overview of the pipeline used to generate cytokine-specific binders. On top of the standard “ColabDesign → TriFlow → AlphaFold3” pipeline, we added all-against-all binding evaluation between designed binders and all class I cytokines in humans to evaluate the off-target effects. **b)** Post hoc analysis showing the scaling relationship between predicted binder specificity and the number of binder sequences designed for each target. Specificity is quantified as the difference in ipAE_min_ between the intended target and the best off-target, and this value is the average for 100 random samples of a certain number of designed sequences to minimize the random effects. **c)** Heatmap depicting on-target and off-target interactions for each selected binder (with optimal specificity score) based on ipAE_min_ scores, highlighting the specificity achieved for each cytokine binder. The designed binders are ordered from the left to the right on the X-axis by decreasing specificity score, and their corresponding targets are ordered from the top to the bottom on the Y-axis. **d)** Representative AlphaFold3 prediction of designed binders targeting IL15, IL21 and IL23A, which shows a low on-target ipAE_min_ (blue, diagonal), poor off-target interaction with the other two targets indicated by a high ipAE_min_, and unfavorable structure prediction showcased by low predicted TM-score for the binder (grey, off-diagonal).

We designed between 4 and 241 sequences per target that satisfied the AlphaFold3-based criteria for on-target binding (Supplementary **Fig. S12c**). After selecting the binder with the highest specificity score for each target, we evaluated its predicted interactions with every class I cytokine using ipAE_min_ (**Fig. 5c**). Although the optimization procedure explicitly maximizes the difference between on-target and off-target ipAE_min_ values, the selected binders also exhibit a clear separation between on-target and off-target ipTM scores (Supplementary **Fig. S12b**). Representative examples for IL15, IL21, and IL23A demonstrate that each binder is predicted to bind its intended target with high confidence while exhibiting low-confidence interactions with the other cytokines (**Fig. 5d**). In several off-target predictions, such as the IL15 binder with IL23A and the IL23A binder with IL21, the binder also fails to maintain its designed structure, further supporting the absence of productive off-target binding.

### Designed sequence profiles informing functional site identification

Evolutionary constraints, which can be inferred from multiple sequence alignments (MSA) across natural protein homologs, have long been essential for inferring protein structure and function^34^. Because TriFlow can generate diverse sequences conditioned solely on a target structure, we sought to ask whether it could help disentangle the contributions of structural versus functional constraints in protein evolution. To this end, we designed sequences for 43,622 diverse protein domains spanning all major evolutionary groups represented in the Evolutionary Classification of Domains (ECOD) database^11^.

We compared conservation patterns in natural sequence profiles with those in the TriFlow-designed profiles for the same domains (**Fig. 6a**). The largest discrepancies occur at functional sites, particularly active sites (mainly catalytic enzyme residues) and small-molecule binding sites annotated in UniProt^35^. These positions are under strong purifying selection in evolution and therefore highly conserved in natural sequences, whereas such functional constraints are absent in structure-conditioned sequence design^36^. Accordingly, designed sequences show only moderate conservation at small-molecule binding residues and almost no conservation at active sites. This likely reflects the fact that small-molecule binding pockets often require unusual or precise structural features, whereas catalytic residues impart relatively little constraint on the 3D structures. Interestingly, post-translational modification (PTM) sites are poorly conserved in both natural and designed profiles, consistent with the observation that PTM positions are not strongly conserved across species^37^.

**Figure 6.**
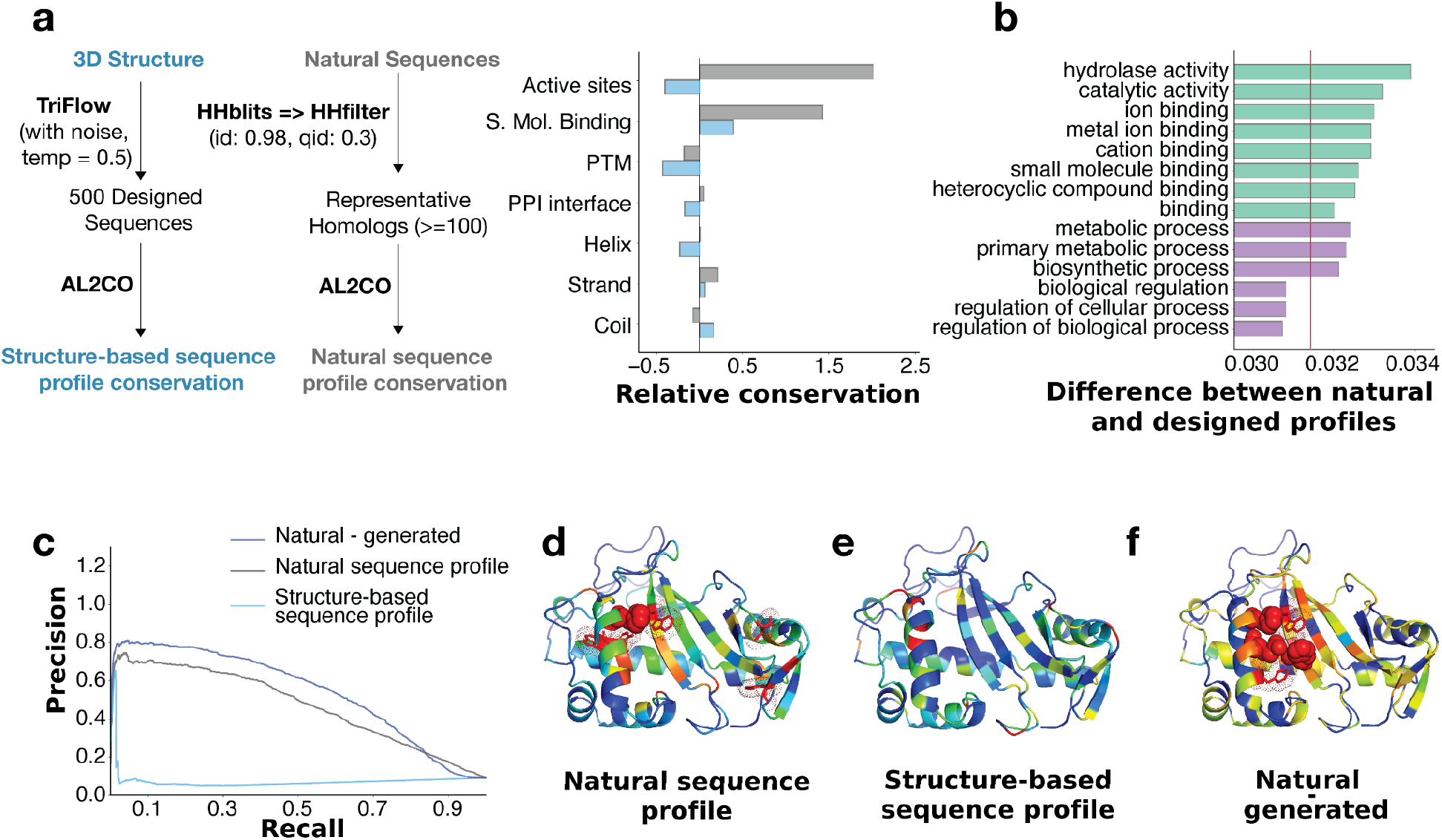
Functional site prediction with TriFlow-designed profiles. **a)** A pipeline to generate designed (left) and natural (mid) sequence profiles and compute sequence conservation. The right plot shows the relative conservation (compared to the average over all residues) of different residue types in natural and designed sequence profiles. **b)** Common Gene Ontology (GO) terms (>1,000 proteins) showing significant differences between natural and designed profiles. Such differences are quantified by Wasserstein Distance at each position, averaged across proteins. The red line indicates the average distance between natural and designed profile over all positions in all proteins. Molecular functions are colored in green and biological processes are colored in purple. **c)** Precision and recall curves at a positive-to-negative ratio of 1:10 for functional site (active site and small molecule binding site) prediction based on sequence conservation in natural profile, designed profile, or the difference between natural and designed profiles. **d-f)** 3D structure of a protein colored by conservation in **d)** natural profile, **e)** designed profile, **f)** the difference between natural and designed profile.

In contrast to functional sites, loops and coils are more conserved in designed sequences than in natural sequences. In natural proteins, these regions are structurally flexible and tolerate extensive sequence variation^38^. However, because current design pipelines receive a fixed backbone as input, even in loops and coils, these positions may experience artificially strong structural constraints that lead to elevated conservation in designed profiles.

Consistent with these residue-level observations, we also found that at the whole-protein level, enzymes and small-molecule–binding proteins show the largest divergence between natural and designed profiles (**Fig. 6b**). In contrast, proteins that function primarily through protein-protein interactions (PPIs), such as regulatory proteins, display relatively small differences between natural and designed sequences.

The pronounced differences between natural and designed sequence profiles at functional sites suggest a new strategy for identifying residues under strong functional constraints. By subtracting the conservation in designed sequences from that in natural sequences, we can effectively remove the contribution of structural constraints and highlight positions that are intolerant to mutation due to functional constraints. This simple approach markedly improves the precision–recall relationship for distinguishing true functional sites from background positions based on sequence conservation (blue versus grey curves in **Fig. 6c**).

An illustrative example is the fungal ribonuclease Rh, whose structure is shown in **Fig. 6d,e** and colored from red to blue according to decreasing sequence conservation. In the natural profile, one of the five most conserved residues corresponds to a catalytic site (red sphere), and another lies adjacent to this active residue facing the substrate-binding pocket; the remaining highly conserved positions likely reflect structural constraints. In contrast, the most conserved residues in the designed profile track unusual structural features and frequently occur in flexible loops. Notably, comparing the conservation patterns of natural and designed sequences highlights functional residues more effectively than conservation in natural sequences alone. Positions with the largest conservation differences correspond either to active-site residues (red spheres in **Fig. 6f**) or to neighboring residues lining the substrate-binding groove (red sticks in **Fig. 6f**).

## Discussion

We presented a systematic study of structure-conditioned sequence design space that spans generative methodology, model architecture, training data expansion, large-scale benchmarks, and biomedical applications. TriFlow advances structure-conditioned protein sequence design by unifying a global three-track RoseTTAFold/AlphaFold-like architecture with discrete flow matching. In contrast to autoregressive MPNN-based approaches, TriFlow leverages global structural context and parallel decoding to capture long-range dependencies across the entire protein or complex, enabling efficient few-step sequence generation for backbones of any length. Our results show that this architectural shift leads to consistent gains in both monomer and binder design performance.

In the monomer design setting, TriFlow outperforms existing sequence-design methods in both native sequence recovery and refoldability. The advantage is particularly pronounced for large backbones, where long-range structural constraints are critical. Introducing backbone noise during training reduces exact sequence recovery but improves refoldability, suggesting that TriFlow learns to produce sequences that are robust to local structural perturbations rather than memorizing native residue identities. The use of Evoformer-like pair representations helps TriFlow to learn inter-residue dependencies and use it to guide sequence generation. TriFlow-designed MSAs recover residue-residue couplings that align more closely with 3D contacts than those generated by other tools. Together, these findings indicate that TriFlow learns a sequence distribution that encodes global structural constraints more faithfully than prior models.

A major motivation for developing TriFlow was to improve *de novo* binder design. Across both a curated set of therapeutic targets and a new large-scale benchmark, TriFlow consistently outperforms ProteinMPNN and SolubleMPNN as a drop-in replacement in state-of-the-art pipelines based on RFdiffusion and BindCraft. In the BindCraft pipeline, SolubleMPNN was only used to redesign the non-interface residues because keeping the interface residues designed by ColabDesign increases the pipeline’s success rate. In contrast, TriFlow improves structure success rates regardless of whether the interface residues designed by ColabDesign are retained. This suggests that TriFlow can match the interface optimization achieved by backpropagation through AlphaFold2, despite being trained as a standalone sequence generation model. One plausible explanation is that TriFlow’s AlphaFold-inspired architecture enables it to internalize global structural priors similar to those used by AlphaFold, producing sequences that are intrinsically favored by AlphaFold3. Yet, as a generative model, TriFlow can design diverse sequences much more efficiently than backpropagation through AlphaFold.

It is important to note that our *in silico* evaluations rely on AlphaFold-based metrics as surrogates for experimental success. To assess whether TriFlow can generate functional binders, we performed limited experimental validation, using BindCraft as a reference for benchmarking. Approximately 20% of the designs selected by the *in silico* filters showed evidence of target engagement, comparable to the success rate achieved by the BindCraft pipeline. Although several SPR sensorgrams exhibited non-ideal kinetic behavior, including incomplete saturation during association and deviations from a simple 1:1 binding model, which precluded rigorous affinity estimation, the observed binding responses are nevertheless consistent with target engagement and suggest that TriFlow performs comparably to BindCraft.

In addition to the discrete flow and global attention-based model design, we expanded the training dataset to include interacting domain pairs from predicted structures in AFDB alongside experimentally determined PDB structures. Because inter-domain interfaces share many structural and physicochemical properties with inter-protein interfaces^39^, this strategy exposes the model to a broader diversity of protein interfaces. We found that this expanded dataset has the potential of improving model performance, but also observed that inaccuracies in AlphaFold-predicted structures can have adverse effects. For example, when backbone noise is not incorporated during training, adding AFDB models leads to a slight decrease in performance, likely reflecting errors or systematic biases in the predicted structures.

For robust benchmarking of computational binder design pipelines, we introduced a large and diverse 500-target binder design benchmark (LBD). LBD is more than an order of magnitude larger than prior benchmarks, and we hope it will facilitate better evaluation of binder design methods. Using this benchmark, we systematically analyzed which target and interface properties predict binder design success rate, which provide guidance to applications and directions for future method development. We found that target interfaces made of alpha helices and depleted of positively charged residues tend to represent easier design cases for modern protein-design pipelines. For targets whose interface properties are associated with lower success rates, generating larger numbers of designs can help compensate for reduced performance. We also confirmed that engaging the native binding interface on the target increases design success. However, we observed a lower success rate for targets with large native binding interfaces, which suggests that the size of the binder should scale with the size of the native interface to fully occupy the native binding surface that has been evolutionarily optimized for PPIs.

We found that target interfaces enriched in α-helices and depleted of positively charged residues are generally more amenable to current protein binder design pipelines. For targets with less favorable interface properties, generating larger numbers of designs can partially compensate for lower success rates. We also confirmed that targeting the native binding interface improves design success. However, success rates decrease for targets with large native interfaces, possibly because small designed binders cannot fully engage binding surfaces that have been evolutionarily optimized for binding proteins. This observation suggests that binder size should scale with the extent of the native interface to maximize design success.

Specificity is an important yet understudied dimension of *de novo* binder design. In our design campaign targeting all class I cytokines, we found that although most binders exhibit strong predicted preference for their intended targets, off-target interactions remain non-negligible. We showed that specificity can potentially be improved by increasing the number of designed sequences and explicitly evaluating cross-reactivity against potential off-targets with sequence and structural similarity to the intended target. We provide computationally predicted specific binders for most class I cytokines as a public resource. These designs offer a valuable starting point for experimental validation and illustrate how efficient sequence generators like TriFlow can accelerate the development of highly specific protein binders.

By systematically comparing structure-conditioned sequence profiles with those derived from natural homologs, we found that designed proteins exhibit markedly reduced conservation at enzyme active sites and small-molecule binding pockets. This observation suggests that many catalytic and ligand-binding residues are constrained primarily by function rather than by the structural requirements of the global fold, and are therefore not recovered by sequence design methods that optimize only for structural compatibility. Subtracting conservation in designed sequences from natural conservation enhances the identification of functional residues relative to natural sequence conservation alone, highlighting the potential of structure-conditioned sequence design to aid functional annotation and improve our understanding of protein function.

Finally, our analyses also highlight several limitations and corresponding opportunities for advancing TriFlow and structure-conditioned sequence design. First, current models optimize sequences solely for structural compatibility, without incorporating functional requirements; integrating explicit functional objectives, such as catalytic efficiency or ligand specificity, into training or sampling could enable true joint structure-function design. Second, existing pipelines assume rigid backbones even in loops and coils, producing artificially high conservation in flexible regions; incorporating backbone ensembles, conformational dynamics, or explicit flexibility during design would better capture natural sequence variation. Third, sequence design and structure generation are currently performed separately, limiting the ability to co-optimize both components; coupling TriFlow with generative structure models offers a way to jointly refine backbone and sequence for improved protein design performance. Finally, the designed binders produced by established pipelines are overwhelmingly α-helical, whereas natural PPI interfaces frequently rely on diverse structural motifs, including β-strands, mixed secondary structures, and irregular loops, to achieve high affinity and specificity. This bias may not only arise from the structure-generation tools but also because α-helical scaffolds are more readily folded and confidently predicted by AlphaFold, even without MSAs. Overcoming this limitation will likely require a new generation of structure-prediction and backbone-generation models that can more faithfully capture the diversity of natural folds and PPI interfaces.

In summary, TriFlow provides a general framework for structure-conditioned protein sequence design that is broadly applicable across protein engineering tasks. To facilitate its adoption, we have released an open-source implementation (https://github.com/jzhoulab/TriFlow), together with user-friendly tutorials. We have also made available the designed sequences for diverse protein backbones and the designed binders targeting diverse proteins (triflow.zhoulab.io), providing a resource for future protein engineering and computational design studies.

## Materials and Methods

### Model Architecture

The TriFlow architecture follows the design of modern protein structure predictors, where multiple iterative attention-based blocks update single, pair, and 3D rigid frames of residues in a protein that exchange information across representations. Single representations are updated using gated attention with pair bias, allowing pair features to modulate attention weights. 3D coordinate updates combine invariant point attention with backbone transformations to maintain rotational and translational consistency. Pair representations are refined through triangle multiplicative updates, like in AlphaFold’s Evoformer, to capture higher-order residue interactions.

Before the main TriFlow blocks, we run a single triangle block comprising triangle multiplicative updates and triangle attention. We use triangle attention in addition to triangle multiplicative update, as it has been shown to be important, especially for structure prediction, in constructing dense 2D distance embeddings. Because triangle attention is expensive and the backbone does not change across timesteps, we compute the triangle block once at the start, cache the resulting pair embeddings, and reuse them for all sampling steps. This reduces computational cost by running the triangle block once rather than running it independently at all 10 sampling steps.

### Training

We trained a version of TriFlow on the same dataset as ProteinMPNN, which are protein assemblies in the PDB until August 2021, clustered at 30% sequence identity and split into train, validation and test sets, ensuring assemblies belonging to different sets never share proteins from the same cluster. We additionally trained a version of TriFlow with additional representative entries from AFDB and predicted domain-domain interactions^12^ from AFDB models to enrich interface design.

The model was initially trained with a crop size of 256 residues and later with 512 residues, using spatial crops centered at the interface in 50% of the samples and contiguous segments in the remaining samples. During training, the backbone input was left unperturbed in half of the samples, while in the other half it was corrupted with Gaussian noise (standard deviation = 0.2 Å) to improve robustness. In addition, the model is provided with a noise token, which encodes if the backbones are perturbed. We trained the final TriFlow model on AFDB using two backbone noise settings. Half of all training steps used a low noise level of 0.02 Å, and the remaining half used a higher noise level of 0.2 Å. At each step, the backbone was perturbed with one of these levels sampled with equal probability.

Training followed a discrete flow-matching objective that linearly interpolates between the masked state, an extra state beyond the 20 canonical amino acids, and the native sequence along a continuous-time Markov chain trajectory. We specifically follow discrete flow models implementation for training and sampling (Supplementary **Fig. S13** and **Fig. S14**). The model was optimized using cross-entropy loss between the predicted logits and the true one-hot-encoded amino acid sequence. We also include a distogram auxiliary loss, similar to AlphaFold2, on the backbone atoms.

We use the AdamW optimizer with a learning rate warmup of 1000 steps from 0 to 0.001. We train the model with a batch size of 8 per GPU on 4 A100 GPUs. For the TriFlow model trained on both PDB and AFDB data, we initialized the model weights with the pre-trained weights from the model trained on ProteinMPNN training dataset.

### Sequence Recovery, Refoldability and Residue-Residue Interaction Analysis

To evaluate sequence recovery, we filtered the ProteinMPNN test set to include only PDB biological assemblies with fewer than 1000 residues and randomly sampled one assembly per cluster. We compared TriFlow trained on the ProteinMPNN training set, ProteinMPNN (v_48_002.pt model weight was chosen because it had the best sequence recovery), ESM-IF and PiFold. To assemble the refoldability benchmark, we used the pregenerated structures with relaxed sequence optimization (RSO) in ColabDesign from a previous study.^13^ We sampled eight sequences per structure and compared TriFlow, trained on the ProteinMPNN dataset with 0.2 Å backbone noise, against ProteinMPNN (trained with 0.2 Å backbone noise), ESM-IF, and PiFold. For all models, we used a sampling temperature of 0.1, except for PiFold, for which we always picked the amino acid with maximal probability following the authors’ recommendation. We used AlphaFold3 in single-sequence mode, without MSAs or templates as inputs.

We assessed whether the backbone-conditioned sequence design distribution reflected explicit structural constraints and whether the model learned dependencies (coevolution) between different positions. We randomly selected 30 biological assemblies (single PDB chains) from the ProteinMPNN test set. We generated 1000 sequences for each biological assembly by three methods, including TriFlow, ProteinMPNN trained on 0.2 Å noise, and ESM-IF, to create the generated profiles. To check whether the generated profiles contained information about structural constraints, we used GREMLIN to identify coevolving residues. We evaluated sampling temperatures ranging from 0.1 to 1.0 for all methods and, for TriFlow, additionally evaluated designs using backbones with or without added noise.

### Binder Design Benchmarks

For the RFdiffusion benchmark, we curated target structures from prior studies (RFdiffusion and a previous Rosetta-based binder design work) and generated 200 structures with RFdiffusion. We designed eight sequences per backbone with TriFlow and ProteinMPNN, respectively. Hotspots were selected from the literature when available; otherwise, they were randomly sampled from the binding interface targeted by previously successful de novo binders. We ran the BindCraft pipeline on its original benchmark set using the interface hotspots specified by the authors (**Table S1**). For each target, the pipeline was run for 2 days on a single A100 GPU, and all generated backbone structures were included in our benchmarking.

A binder sequence was considered successful *in silico* if it met established criteria derived from AlphaFold2 and AlphaFold3 evaluations of the target-binder complex^40^. We followed the AlphaFold2 implementation with initial guess derived from the target structure and without MSAs (single-sequence mode), and used the following cutoffs from the AlphaProteo^40^ benchmarks to select successful binders: binder RMSD < 1.5, average pAE of interface residues < 7, and binder pLDDT > 90. For AlphaFold3, we use the target chain as a template and run the model in single-sequence mode (https://github.com/Kuhlman-Lab/AlphaFold3)^40,41^. We ran three rounds of recycling in AlphaFold3 and used the following cutoffs to select successful binders: ipAE_min_ < 1.5, binder pTM > 0.8, and complex RMSD < 2.5. The AlphaFold3 wrapper we used only supports single-chain template target structures, and naively concatenating multiple target chains did not produce valid predictions. Consequently, no designed binder passed the AlphaFold3 filtering criteria for the multi-chain targets H3 in the RFdiffusion benchmark and Der f21 in the BindCraft benchmark.

### Large-scale Binder Design Benchmarks

We collected 53,471 PDB entries released between August 2021 and April 2025, extracted 71,678 biological assemblies, and identified 955,925 interacting chain pairs in which each chain contained at least five residues within 5 Å of the other chain. To avoid overlap with the training data for ProteinMPNN and TriFlow, we excluded interacting pairs whose close homologs (detected by MMseqs at >30% sequence identity and >60% query or hit coverage) appeared in PDB entries released prior to August 2021. This filtering yielded 297,486 interacting PDB chains. We clustered these chains using MMseqs^42^ (50% identity, 80% coverage) and selected one representative pair per cluster-cluster combination, resulting in 15,367 representative interacting pairs. Most of these pairs originate from higher-order complexes, for which disrupting native assemblies using designed binders might be more challenging. We therefore focused on the 772 representative pairs derived from binary complexes and randomly selected 500 to construct the LBD benchmark. For each binary complex, we designated chain A as the target and used the native binding interface to guide hotspot selection for the RFdiffusion-based pipeline.

We generated binder backbones using both RFdiffusion and ColabDesign from the BindCraft pipeline. To generate binders with RFdiffusion, we provided the native hotspot and generated lengths ranging from 60 to 120 residues. The BindCraft hallucination protocol is computationally expensive, requiring 140 AlphaFold forward passes to produce a single successful trajectory, but a large fraction of generated backbones are discarded by BindCraft’s internal filters, which evaluate structural quality and target engagement. To save time and reduce compute cost, we only took the ColabDesign-based structure generation modules from BindCraft and modified the script to generate structures up to a certain number of trials (specified by a user), rather than the number of successfully generated structures that pass filters. BindCraft would terminate unpromising trajectories if the binder pLDDT went below 65, and thus the number of generated structures could be below the number of trials. This makes the compute time spent on structure generation more predictable and easier to benchmark. We used this ColabDesign script without any hotspots to generate up to 10 backbone structures for each target, requiring each backbone to contain 60-120 residues.

For each backbone generated by RFdiffusion, we performed sequence design with ProteinMPNN and TriFlow, respectively, generating 8 sequences per backbone. For backbones generated by ColabDesign, we used SolubleMPNN from the BindCraft pipeline for comparison against TriFlow. We used TriFlow trained on PDB (prior to August 2021) and AFDB with 0.2 Å noise added to the backbone and set its temperature to 0.1. Similarly, we used ProteinMPNN and SolubleMPNN trained with 0.2 Å noise and set the temperature to 0.1. The BindCraft pipeline by default only redesign the non-interface residues by SolubleMPNN, leaving the interface residues designed by ColaDesign. To test the impact of this strategy, we attempted to redesign all residues or only focus on non-interface residues. We also tried to assume backbone noise (standard deviation of 0.2 Å) in the BindCraft-generated structures and tested whether this assumption affected TriFlow’s performance. During sequence design, cysteine residues were excluded to reduce the risk of unintended disulfide bond formation and associated expression or folding artifacts.

### Design Specific Binders for Human Class I Cytokines

We gathered the predicted 3D structures for 31 human Class I cytokines from AFDB and removed their signal peptides following UniProt’s annotations. We initialized 10 binder backbone generation trials for each cytokine with ColabDesign adapted from BindCraft. However, a fraction of these trials would be automatically terminated due to the pLDDT-based checkpoint (see previous section about our modifications to the BindCraft script), resulting in 2-8 successfully generated backbone structures for each cytokine. We designed 40 sequences using TriFlow trained on both PDB and AFDB for each generated backbone structure. Across the 31 cytokine targets, all but one (IL7) yielded at least one binder passing the same AlphaFold3 thresholds as we used in the binder design benchmarks (ipAE_min_ < 1.5, binder pTM > 0.8, and complex RMSD < 2.5).

To estimate the cross reactivity to other targets, we computed a specificity score, *S_spec_*, for each binder *x* intended to target cytokine *T_t_* based on the following formula:

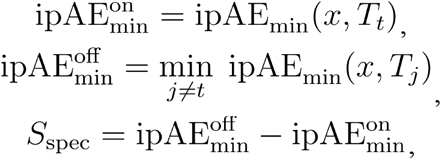

where ipAE_min_ is the minimal interface pAE computed by AlphaFold3 and *T_j_* is any Class I cytokine in our target set. We aimed to identify the binder with the highest *S_spec_*, while satisfying another criterion of binder pTM > 0.8 estimated by AlphaFold3, among our pool of designed binders for each cytokine target. These optimized sequences became our final set of designed specific binders for human class I cytokines (**Fig. 5c**).

We conducted a post hoc analysis to examine the relationship between the maximal binder *S_spec_* for each cytokine target and the number of sequences, *N_design_*, designed for each binder backbone. To examine this relationship, we scanned different values of *N_design_*. For each value of *N_design_*, we randomly sampled *N_design_* designed sequences for each binder backbone and repeated the sampling 100 times to calculate the average maximal *S_spec_* (**Fig. 5b**).

### Comparison of Natural and Designed Profiles and Functional Site Prediction

We clustered ECOD domains parsed from PDB structures using MMseqs (-c 0.8 --min-seq-id 0.4) and selected one representative domain per cluster. For each representative, we used its structure as input to TriFlow trained with 0.2 Å backbone coordinate noise and generated 500 sequences at a sampling temperature of 0.5 to maximize sequence diversity. Because all sequences were generated for the same backbone, they were inherently aligned and collectively constituted the designed sequence profile.

To construct natural MSAs, we searched for homologs of each domain sequence using HHblits^43,44^ against the UniRef30 database. The resulting MSAs were filtered with HHfilter^43,44^ to remove highly similar sequences (sequence identity >98%) and those that were too distantly related to the query (sequence identity <30%). The filtered MSAs were used as the natural sequence profiles. For domains with fewer than 100 sequences remaining after filtering, we omitted the sequence identity filters to retain sufficient sequence depth. Because some domains lacked sufficiently deep MSAs, we restricted downstream analyses to the 19,264 ECOD domains with at least 100 sequences in the MSAs.

We next annotated structural and functional properties for each residue in every ECOD domain. We downloaded UniProt’s reviewed entry annotations (JSON format) and mapped them to the corresponding PDB entries. This allowed us to annotate active sites (catalytic sites), small-molecule binding sites, and PTM sites (including modified residues, glycosylation, and lipidation) in ECOD domains which were derived from PDB entries. In total, 16,976 ECOD domains were mapped to reviewed UniProt entries, allowing us to identify 4,509 active sites, 80,574 small molecule binding sites, and 11,419 PTM sites. Additionally, to identify residues at domain or protein interfaces, we examined the original PDB biological assembly for each domain and labeled residues with any atom within 8 Å of another domain or protein chain as interface residues. Finally, the secondary structure of each residue was analyzed using DSSP^45^.

For each position in every domain, we computed conservation indices for both the natural and designed profiles using AL2CO^46^. For each structural or functional category (as annotated above and shown in **Fig. 6a**), we averaged the conservation indices across all category-associated residues and compared these averages to the global averages across all residues in natural or designed profiles.

To quantify differences between natural and designed profiles at the domain level, we calculated a profile-divergence score for each ECOD domain by averaging the Wasserstein distance^47^ between the natural and designed conservation distributions over all positions. We denote the divergence for domain as *PD_i_* To classify domains into functional categories, we applied ProteInfer^48^ to predict Gene Ontology (GO) terms for each domain. For each GO term, we compared *PD_i_* values of annotated domains with those of domains not annotated with this GO term. Statistical significance was assessed using the Mann–Whitney U test, with false discovery rate (FDR) correction^49^ applied to account for multiple hypothesis testing. We reported GO terms that met the following criteria in **Fig. 6b**: (1) associated with more than 1,000 domains; (2) FDR < 0.05; (3) belonging to the “molecular function” or “biological process” categories.

The marked discrepancy in conservation at active sites between natural and designed profiles prompted us to test whether contrasting these profiles improves functional-site prediction. As positive controls, we used annotated active-site residues in the ECOD domains. Negative controls were all other residues not annotated as active sites, binding sites, or PTM sites in UniProt; the negative set was subsampled to 10× the size of the positive set. We then evaluated whether the two sets could be distinguished using three metrics computed per residue: (1) the AL2CO conservation index computed on the natural profile, (2) the conservation index computed on the designed profile, and (3) the difference between natural and designed conservation indices (**Fig. 6c**).

## Acknowledgments

JZ is supported by the Cancer Prevention and Research Institute of Texas grant RR190071 and by the National Institutes of Health (NIH) grant DP2GM146336. QC is a Southwestern Medical Foundation Endowed Scholar. This research is supported by V-I-0004-20230731 and I-2270-20260402 from the Welch Foundation and R35GM160468 from the National Institutes of Health.

## Supplementary Figures

**Figure S1.**
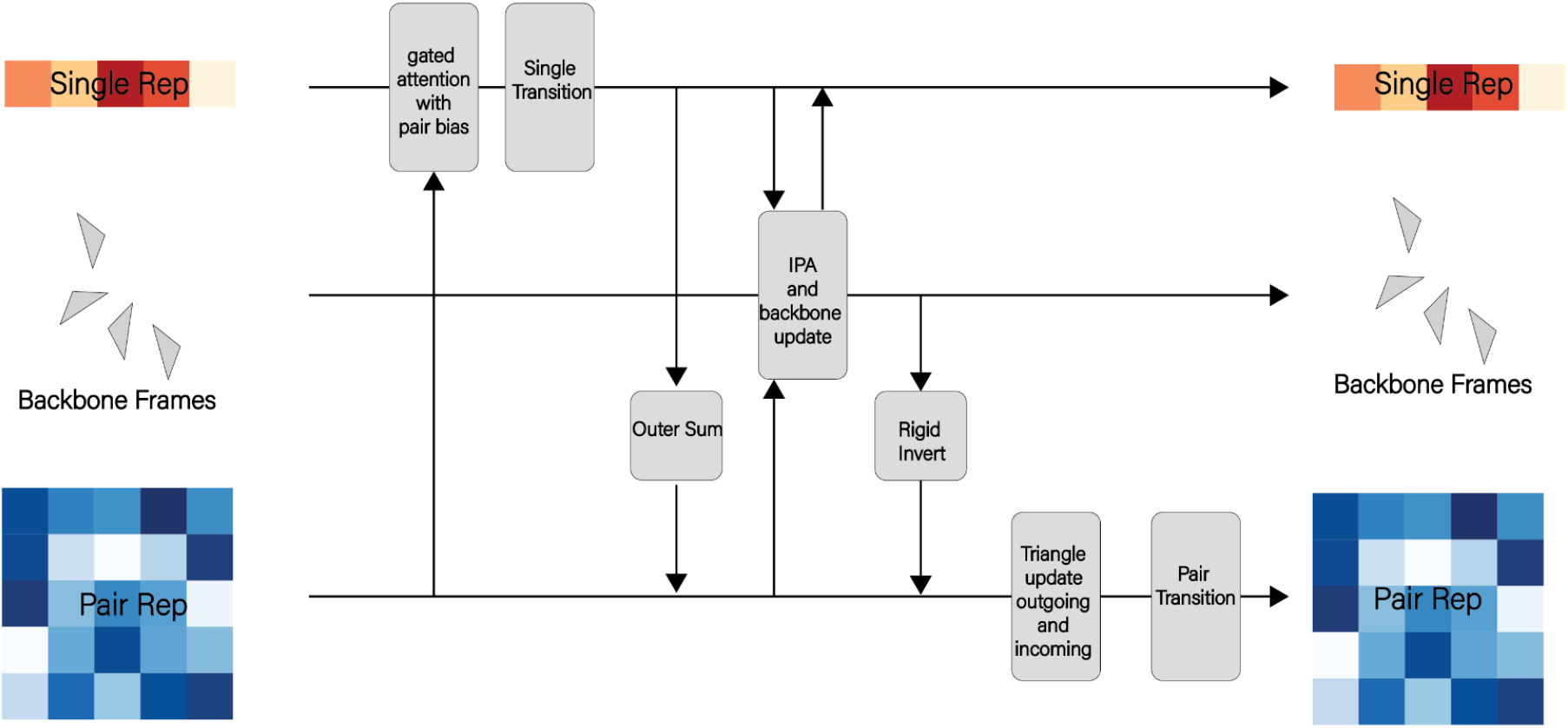
Detailed architecture of TriFlow module showing the information processing and exchange between the three tracks.

**Figure S2.**
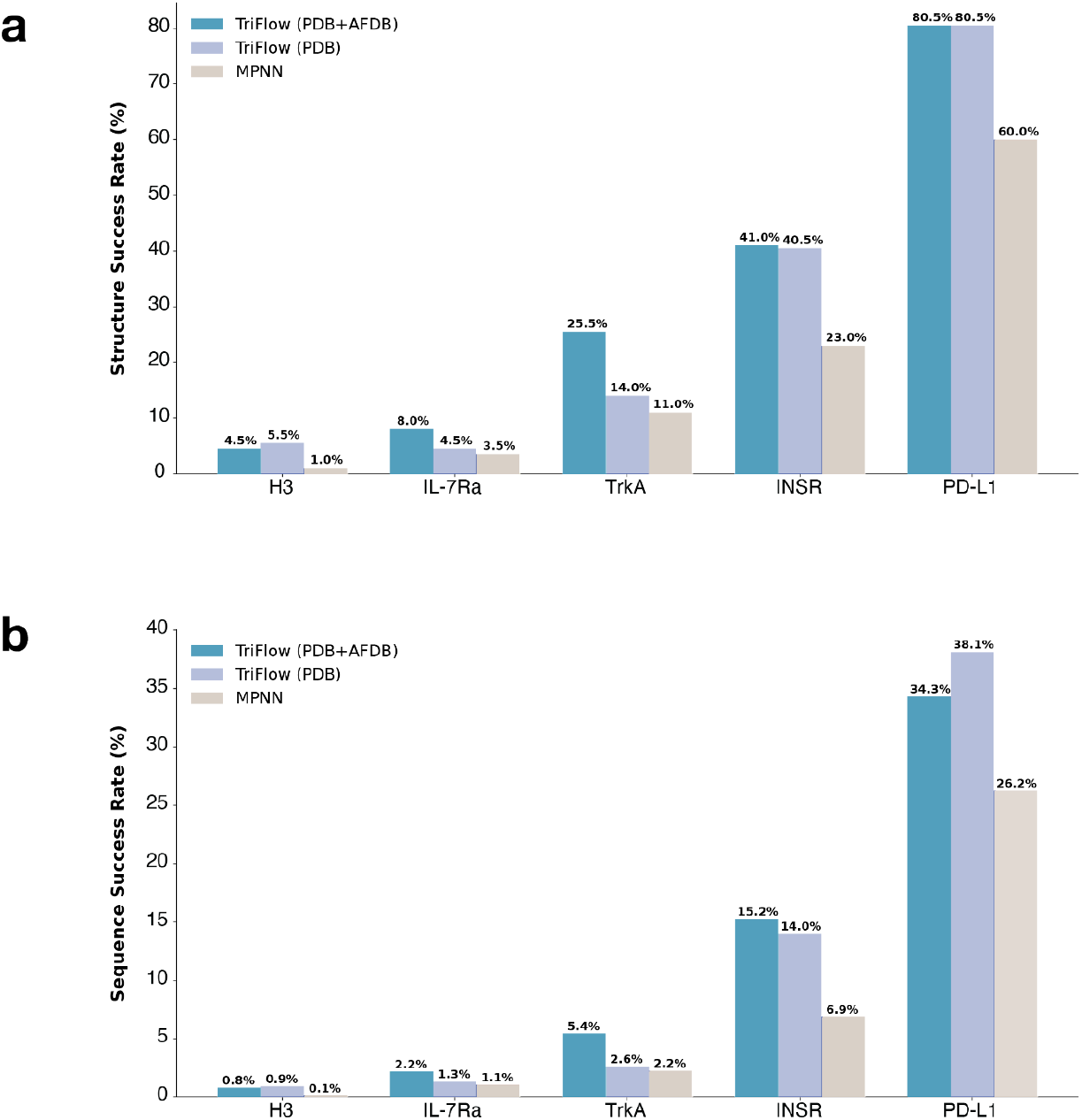
Structure and sequence success rate comparison across training datasets and models. Comparison of **a)** structure and **b)** sequence success rates for TriFlow trained on the PDB dataset, TriFlow trained on both PDB and AFDB datasets, and ProteinMPNN, evaluated on RFdiffusion-generated binder backbones. Structure predictions were performed using AlphaFold2 in single-sequence mode with an initial target template provided.

**Figure S3.**
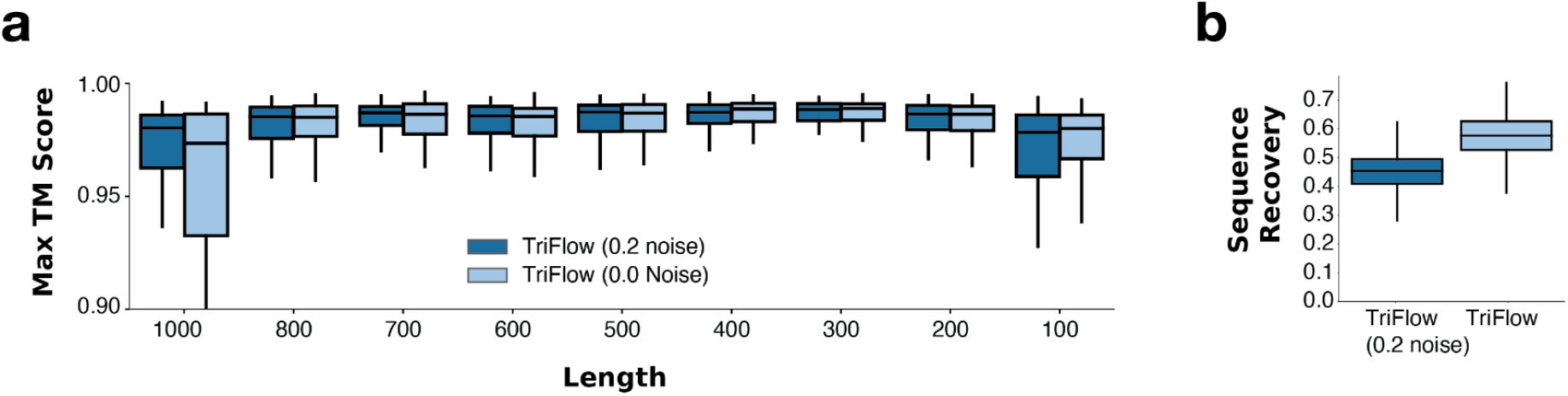
Performance comparison between using noise and not using noise in backbones during sequence generation by TriFlow. **a)** Refoldability of designed sequences for Relaxed sequence optimization (RSO) generated backbone structures; **b)** sequence recovery in the ProteinMPNN test set.

**Figure S4.**
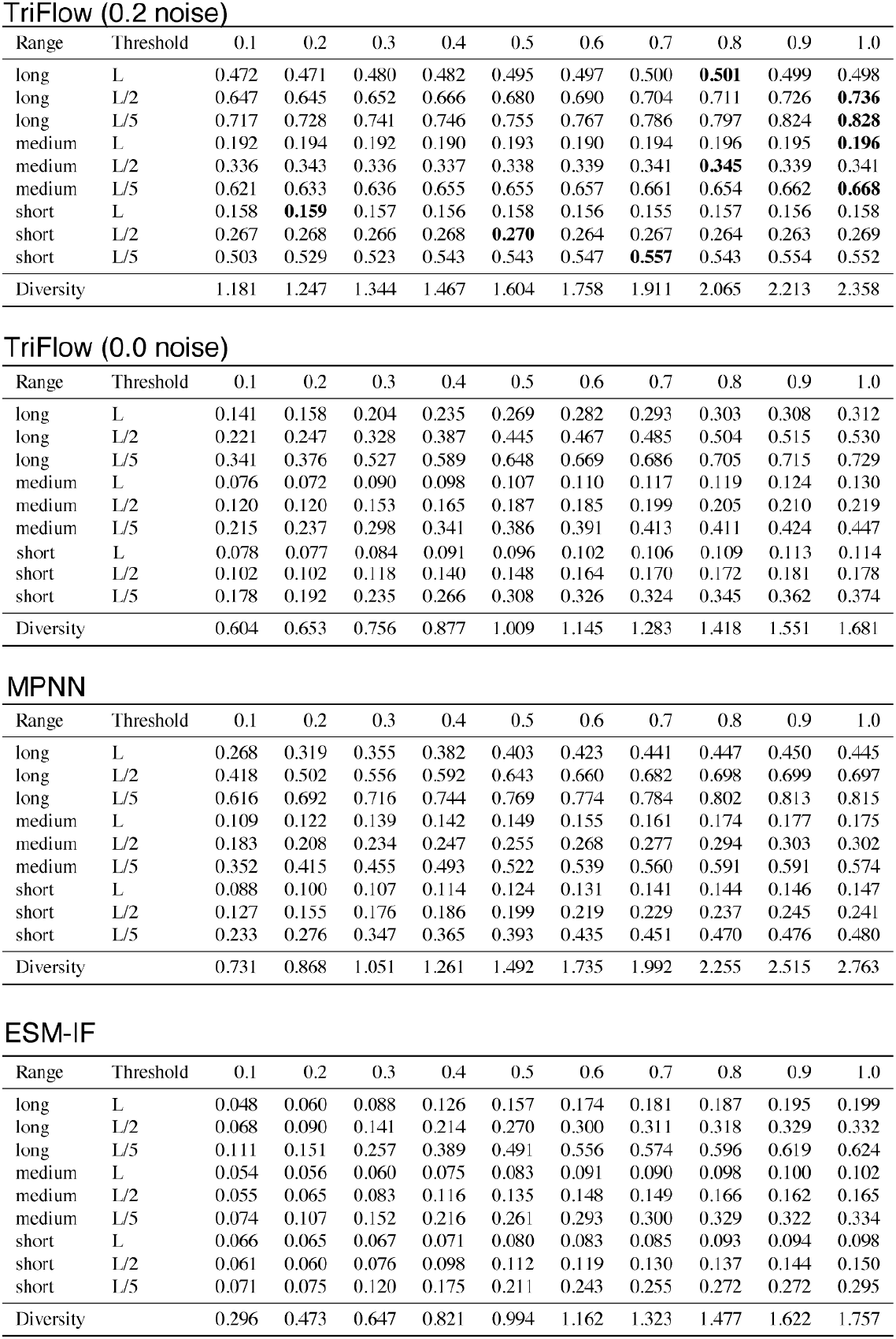
Benchmarking contact prediction accuracy across sequence design models. Contact prediction accuracy was evaluated for TriFlow (with and without backbone noise), ProteinMPNN, and ESM-IF. Precision was computed for the top L, L/2, and L/5 predicted contacts across short [6-12), medium-[12-24), and long-range [>24) sequence separations, as well as across model temperature parameters ranging from 0.1 to 1. Diversity is calculated using Shannon’s entropy on the generated profiles.

**Figure S5.**
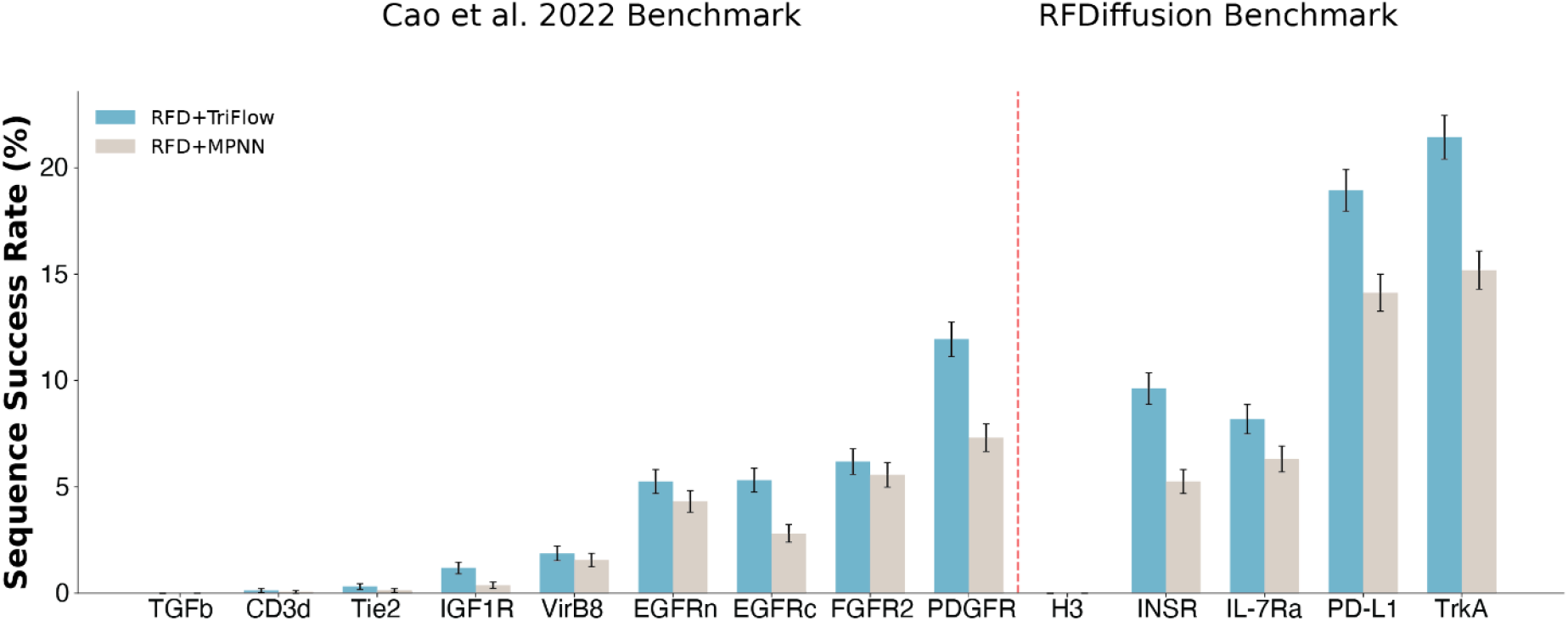
Sequence success rate across target structures. Sequence success rate was defined as the proportion of sequences that passed AlphaFold3 evaluation out of the total 1,600 sequences generated per target. The comparison shows the success rates of sequences generated by TriFlow and ProteinMPNN on backbones produced by RFdiffusion.

**Figure S6.**
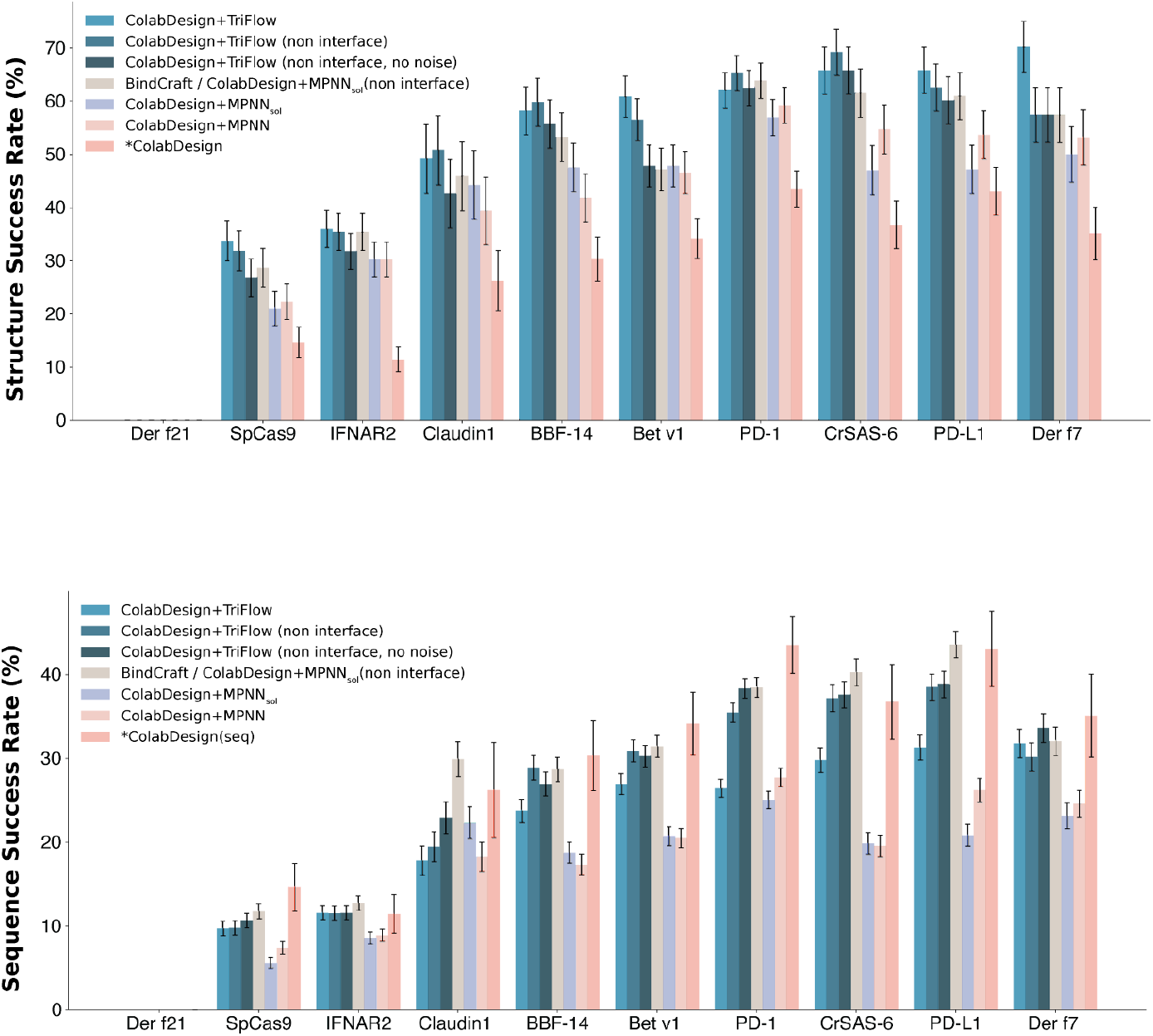
Structure and Sequence success rates for ColabDesign backbones with sequence design model ablations. **a)** Structure success rate comparison across different sequence design strategies. For TriFlow, we evaluated three modes: full binder redesign, redesign restricted to non-interface residues, and sequence design without added structural noise. For ProteinMPNN, analogous ablations were performed, including the BindCraft pipeline (SolubleMPNN redesigning only non-interface residues), full binder redesign with SolubleMPNN, and full redesign with ProteinMPNN. The ColabDesign sequence was obtained directly from backbone generation without any redesign. **b)** Sequence success rates under the same ablation settings for TriFlow and ProteinMPNN.

**Figure S7.**
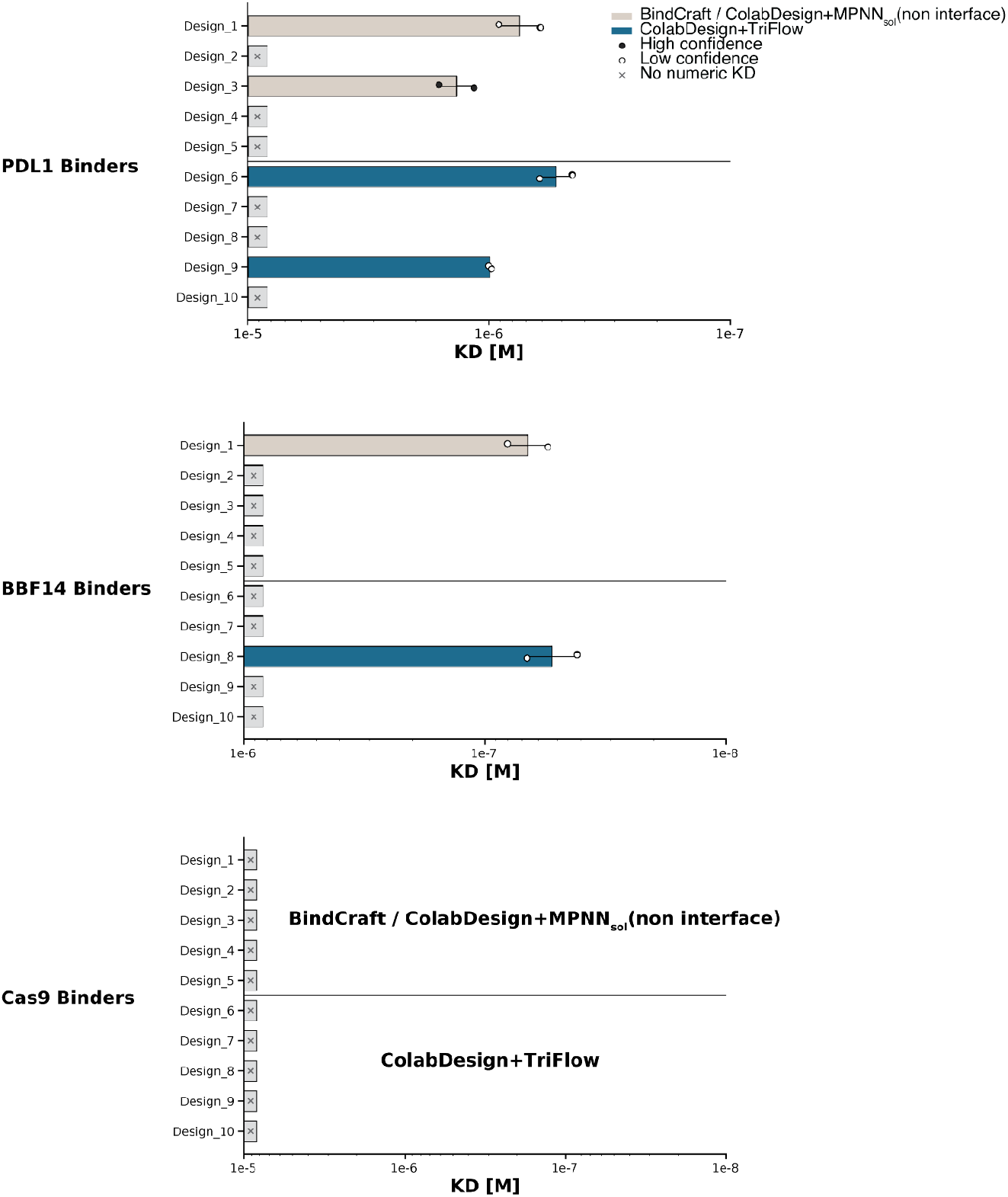
Estimated binding affinity for designed binders of PDL1, BBF14, and Cas9 by the BindCraft pipeline (SolubleMPNN) and TriFlow. For each target, designs 1-5 are by the BindCraft pipeline, designs 6-10 are by TriFlow.

**Figure S8.**
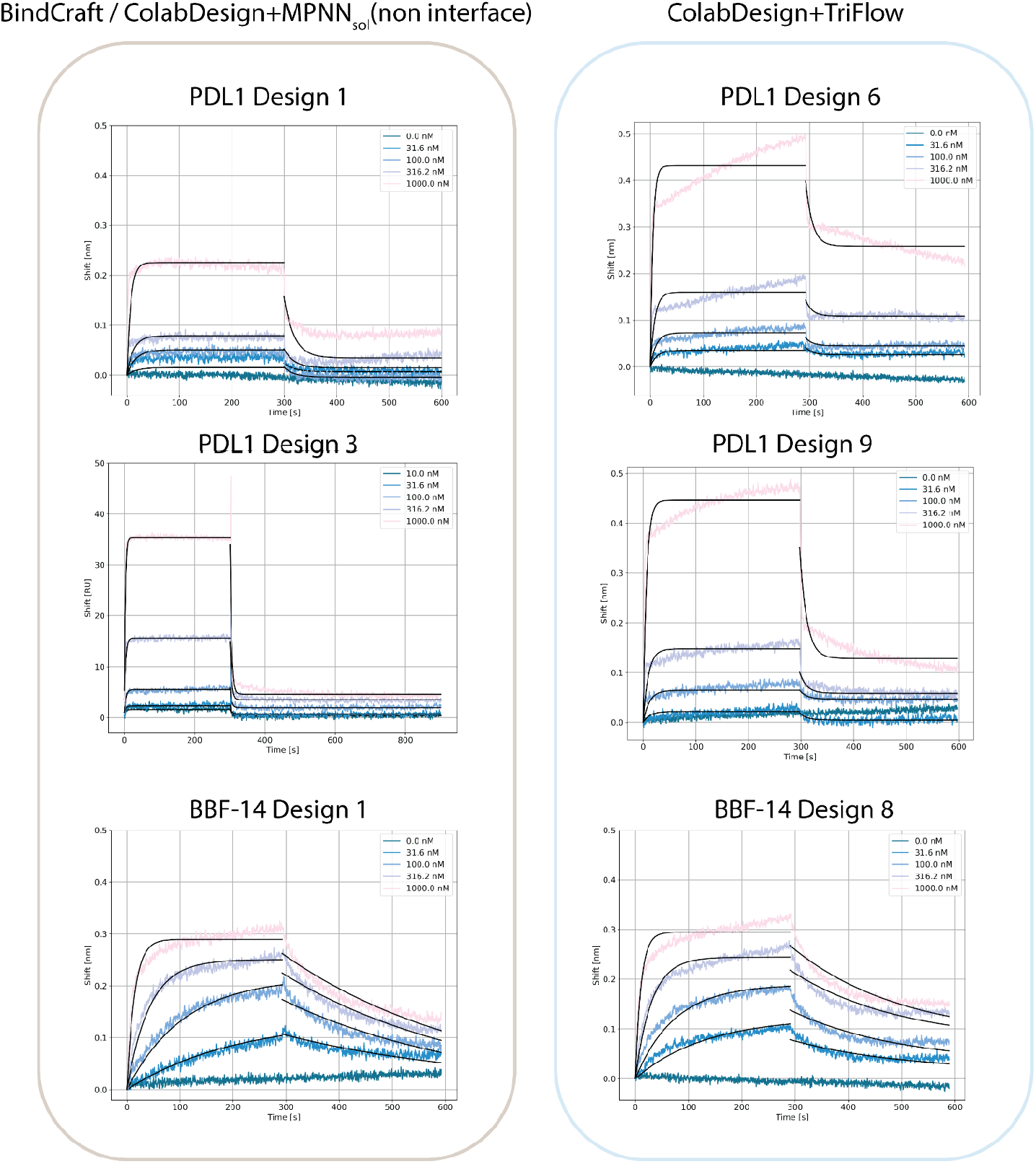
Concentration-dependent BLI and SPR measurements for binders exhibiting detectable binding affinity. Examples of successfully experimentally validated ColabDesign+TriFlow and BindCraft/ColabDesign+MPNN_sol_(non interface).

**Figure S9.**
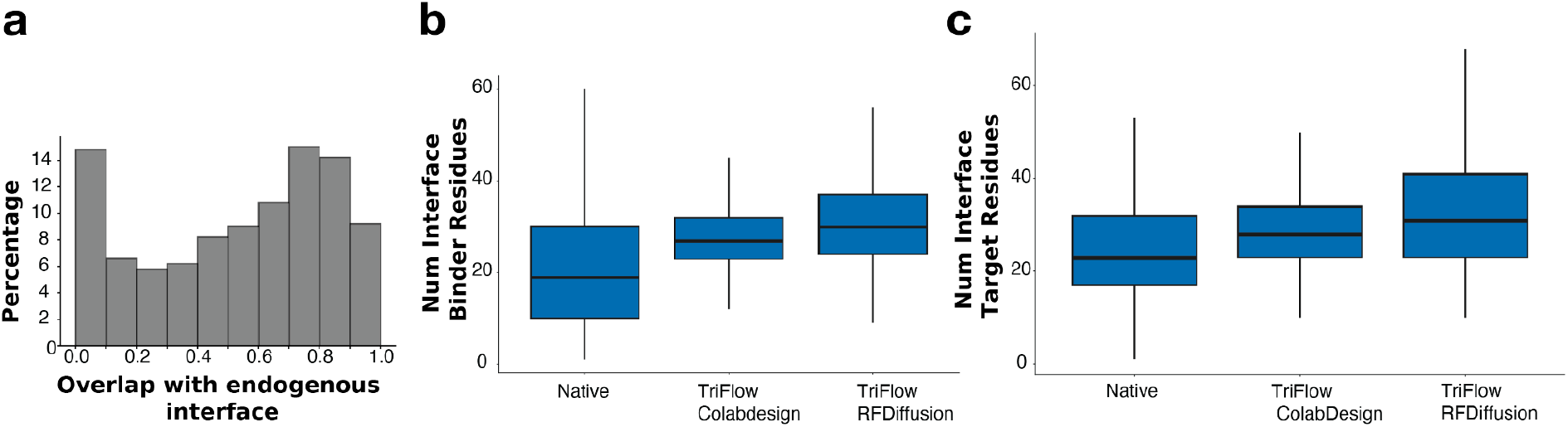
Interface properties of designed and native binders. **a)** Distribution of interface overlap between designed binders by ColabDesign and their corresponding endogenous (native) binders, expressed as the fraction of shared interface residues in ColabDesign binders. **b)** Comparison of the number of interface residues on the binder side across native complexes, TriFlow+ColabDesign binders, and TriFlow+RFdiffusion binders. **c)** Comparison of the number of interface residues on the target side across the same sets of structures.

**Figure S10.**
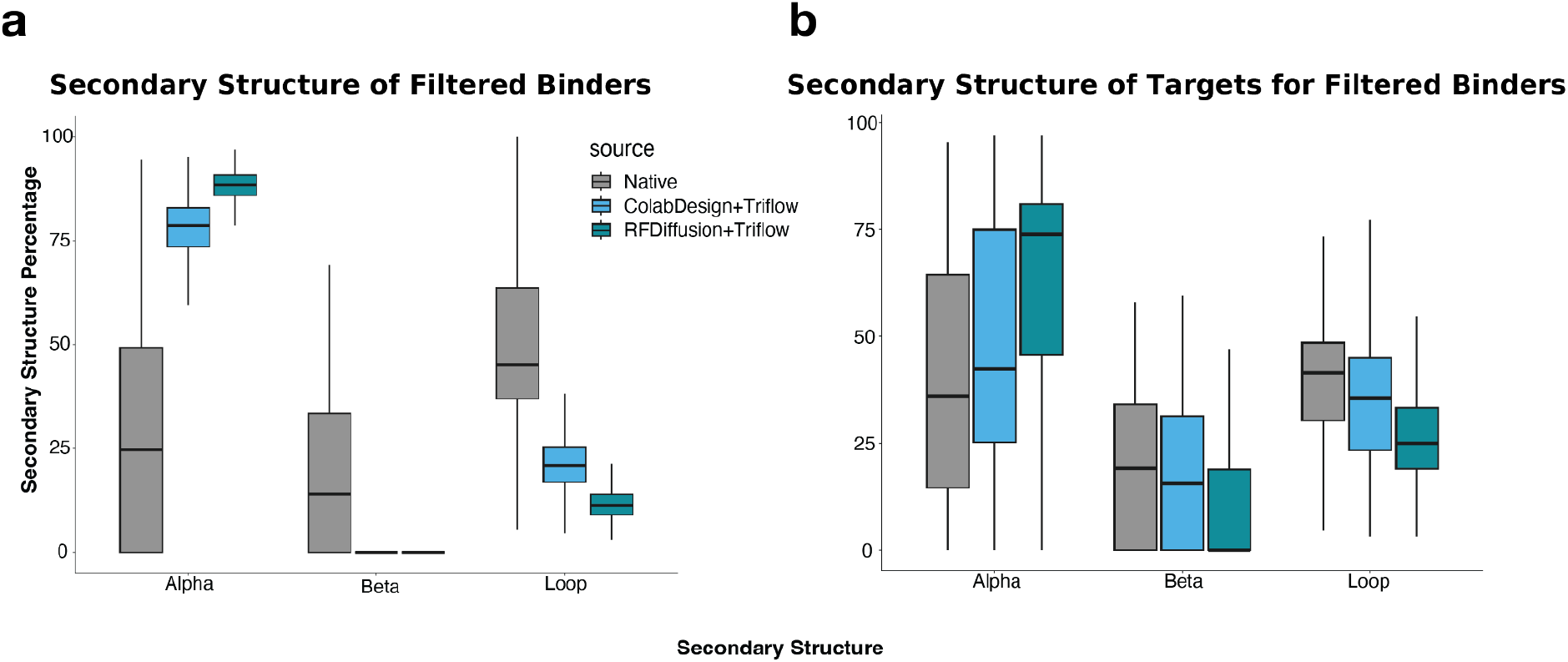
Secondary structure composition of designed binders passing *in silico* filters and their targets. **a)** Comparison of secondary structure content among filtered binders generated by TriFlow with ColabDesign and RFdiffusion backbones, relative to native binders. The analysis shows the proportion of alpha-helical, beta-strand, and loop regions within each binder. **b)** Secondary structure composition of the corresponding target proteins for the same set of filtered binders.

**Figure S11.**
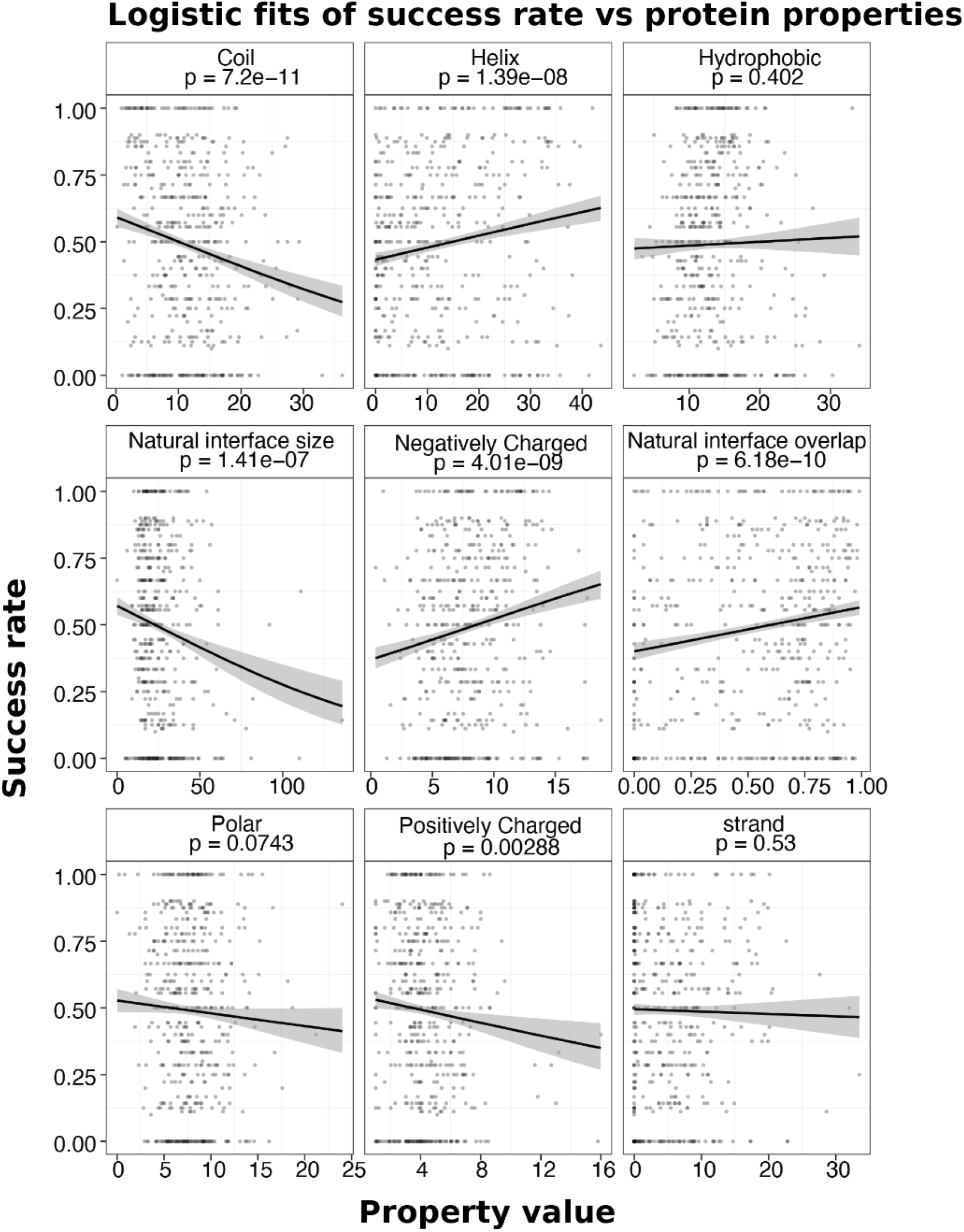
Logistic regression analysis of structural and physical properties influencing design success. Logistic regression models show how both structural and physical features of the target structure correlate with structure success rate. Structural features include secondary structure composition, interface size, and interface overlap between native and designed binders. Physical properties include charge distribution and the balance of hydrophobic and polar groups.

**Figure S12.**
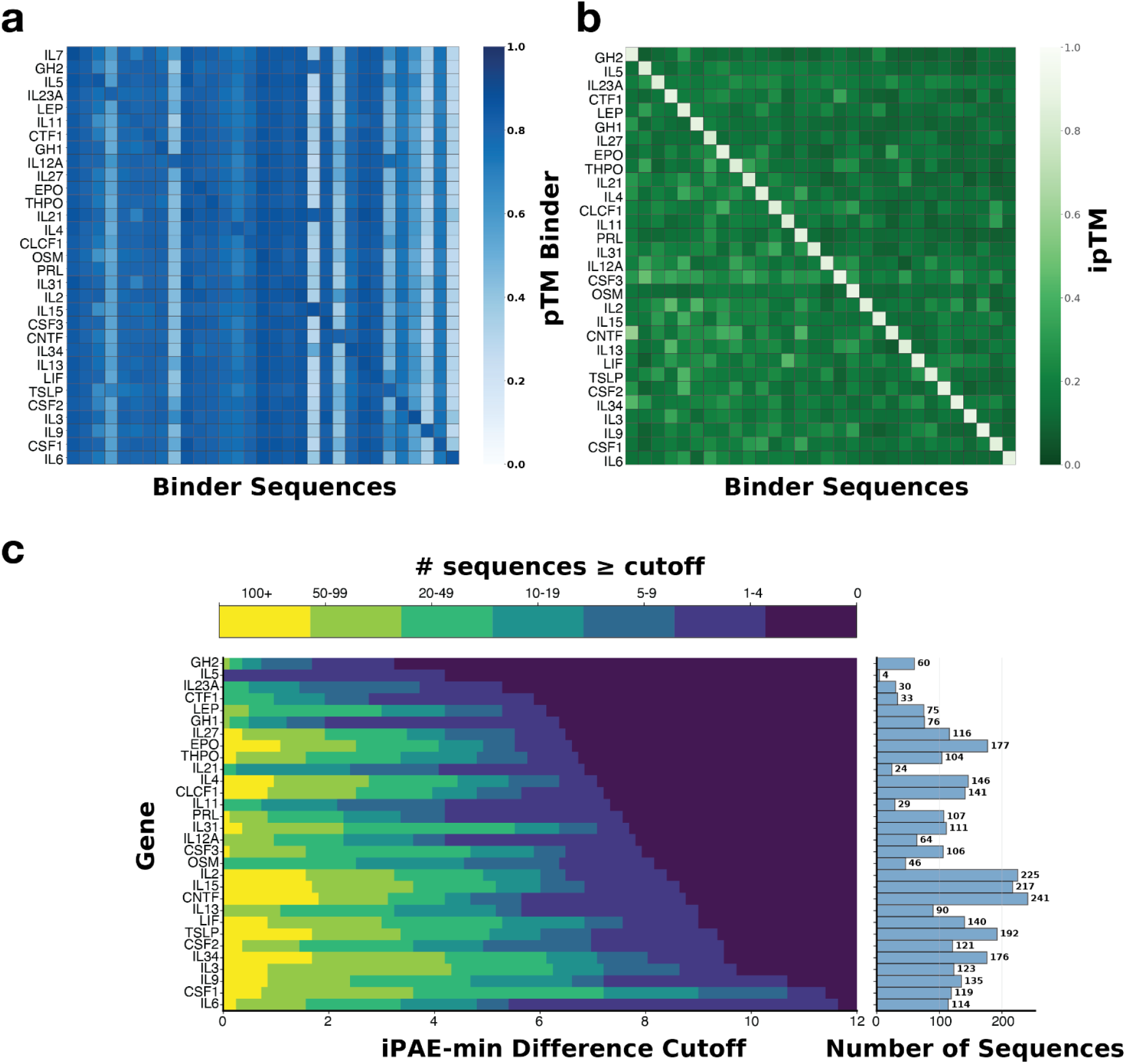
Confidence metrics for generating specific cytokine binders. **a)** Heatmap of predicted TM-scores showing the structural confidence of the best binder for each cytokine target (on-target) and its corresponding off-target predictions against other cytokines. **b)** Comparison of interface predicted TM-scores for on-target versus off-target predictions, demonstrating clear separation between on-target and off-target interactions. **c)** Number of designed sequences available across varying ipAE_min_ difference cutoffs, where the ipAE_min_ difference quantifies the gap between the best on-target ipAE_min_ score and the next-best (lowest) off-target score. The histogram (right) summarizes the total number of designed sequences that pass AlphaFold3 confidence metrics for on-target interactions

**Figure S13.**
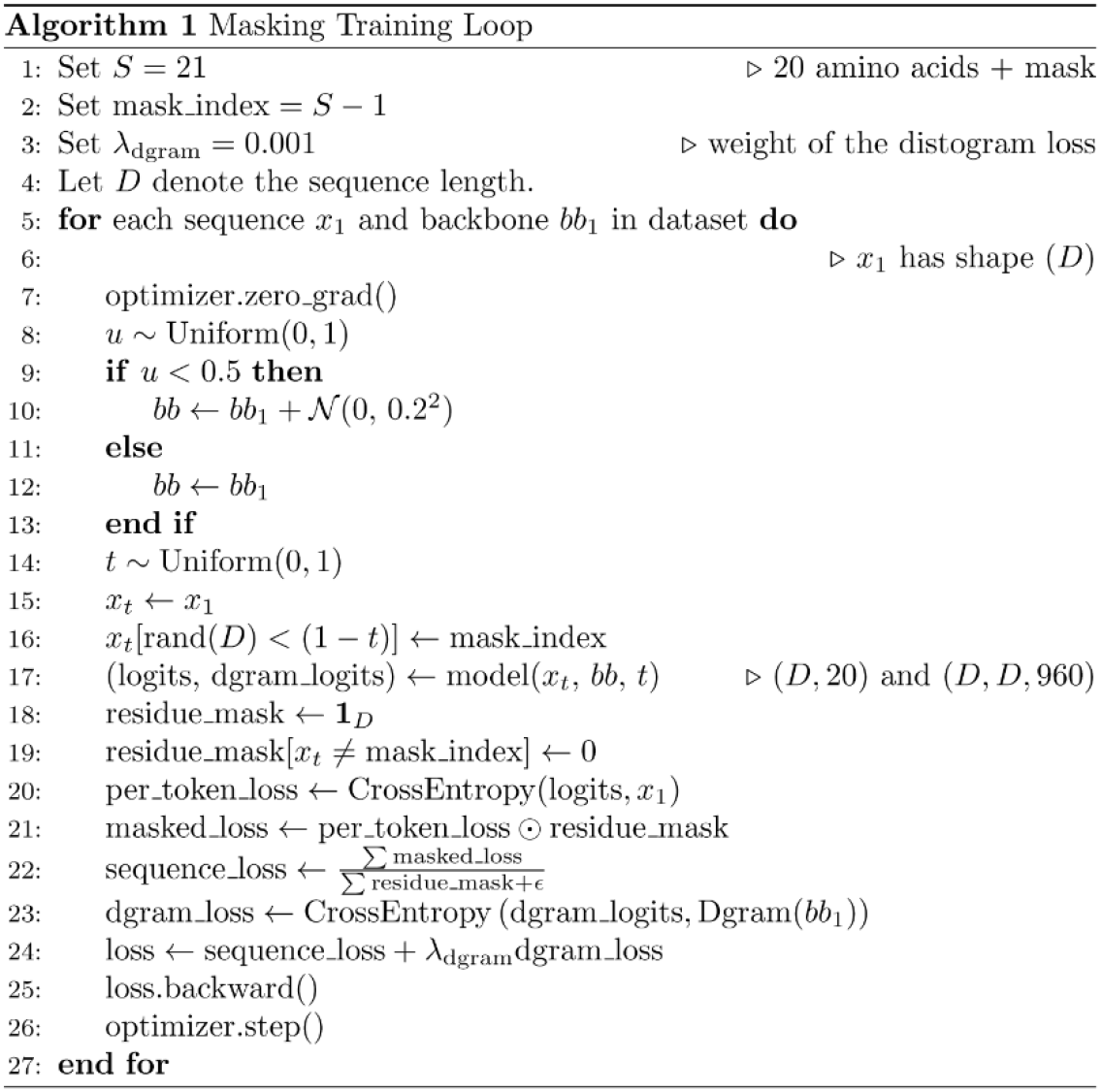
Pseudocode for the masked training loop. During training, random masking based on a scheduler and backbone noise are applied to the input sequence and structure. The model predicts the amino acid identity and the cross entropy loss is calculated over the masked residues.

**Figure S14.**
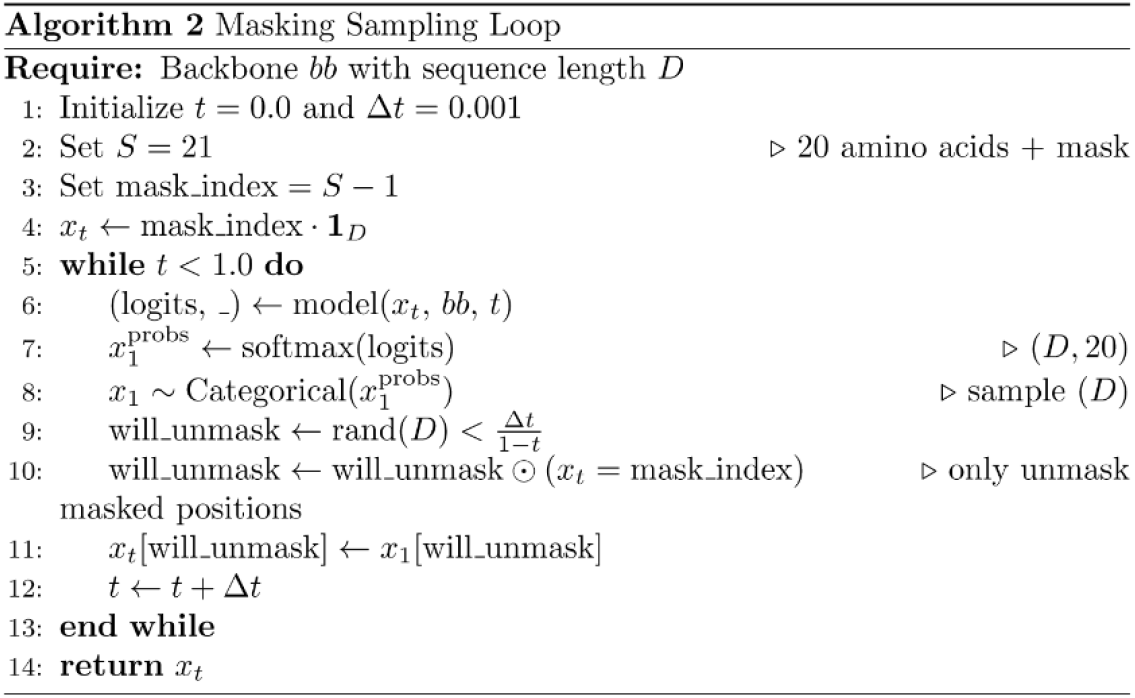
Pseudocode for the inference loop. Inference begins with the sequence initialized entirely as mask tokens. A time-dependent unmasking schedule progressively reveals a subset of the remaining masked residues, whose identities are sampled from the model’s predicted amino acid distributions. This process is repeated until all residues have been assigned an amino acid identity.

**Table S1.** Configuration for running RFdiffusion and BindCraft on the binder design benchmarks.

| Target | Tool | Input PDB | Hotspot | Binder Size |
| --- | --- | --- | --- | --- |
| PD-1 | BindCraft | AF2 prediction, trimmed to 32-146 | 64, 126, 129, 133 | 80-150 |
| PD-L1 | BindCraft | AF2 prediction, trimmed to 18-132 | 54, 56, 66, 115 | 65-155 |
| IFNAR2 | BindCraft | 2LAG, trimmed to 8-110 | 52, 80, 82, 84, 96, 98 | 60-175 |
| Claudin1 | BindCraft | AF2 prediction of CLN1-14 | 31, 46, 55, 152 | 80-175 |
| BBF-14 | BindCraft | 9HAG | none | 70-250 |
| CrSAS-6 | BindCraft | AF2 prediction, trimmed to 15-160 | none | 90-200 |
| Der f7 | BindCraft | AF2 prediction with 3UV1 template, trimmed to 18-213 | 132 | 70-185 |
| Der f21 | BindCraft | AF2 prediction with 5YNY template, trimmed to 25-136 | 34 | 70-185 |
| Bet v1 | BindCraft | AF2 prediction | 24 | 70-185 |
| SpCas9 | BindCraft | AF2 prediction with 4ZT0 template, trimmed to 96-174 + 306-446 | 360 | 70-150 |
| TGFb | RFdiffusion | AF2 prediction | B14,B63,B66,B67 | 40-100 |
| CD3d | RFdiffusion | AF2 prediction | A6,A11,A31,A32 | 40-100 |
| Tie2 | RFdiffusion | AF2 prediction | A127,A137,A138,A171 | 40-100 |
| IGF1R | RFdiffusion | AF2 prediction | A52,A78,A87,A108 | 40-100 |
| Virb8 | RFdiffusion | AF2 prediction | A43,A47,A58,A59 | 40-100 |
| EGFRn | RFdiffusion | AF2 prediction | A8,A9,A95,A99 | 40-100 |
| EGFRc | RFdiffusion | AF2 prediction | A72,A102,A108,A128 | 40-100 |
| FGFR2 | RFdiffusion | AF2 prediction | A4,A48,A87,A89 | 40-100 |
| PDGRF | RFdiffusion | AF2 prediction | A116,A120,A138,A141 | 40-100 |
| H3 | RFdiffusion | 5VLI | B521, B545, B552 | 40-120 |
| INSR | RFdiffusion | 4ZXB | E64, E88, E96 | 40-120 |
| IL-7Ra | RFdiffusion | 3DI3 | B58, B80, B139 | 50-120 |
| PD-L1 | RFdiffusion | 5O45 | A56, A115, A123 | 50-120 |
| TrkA | RFdiffusion | 1WWW | X294, X296, X333 | 50-120 |

## References

1. Pacesa, M. et al. One-shot design of functional protein binders with BindCraft. Nature 646, 483–492 (2025).

2. Watson, J. L. et al. De novo design of protein structure and function with RFdiffusion. Nature 620, 1–3 (07 2023).

3. Jumper, J. et al. Highly accurate protein structure prediction with AlphaFold. Nature 596, 583–589 (2021).

4. Dauparas, J. et al. Robust deep learning–based protein sequence design using ProteinMPNN. Science 378, 49–56 (10 2022).

5. Gao, Z., Tan, C., Chacón, P. & Li, S. Z. PiFold: Toward effective and efficient protein inverse folding. arXiv [cs.AI] (2022).

6. Hsu, C., et al. Learning inverse folding from millions of predicted structures. bioRxiv (2022) doi:10.1101/2022.04.10.487779.

7. Alon, U. & Yahav, E. On the bottleneck of graph neural networks and its practical implications. arXiv [cs.LG] (2020) doi:10.48550/arXiv.2006.05205.

8. Baek, M. et al. Efficient and accurate prediction of protein structure using RoseTTAFold2. bioRxiv 2023.05.24.542179 (2023) doi:10.1101/2023.05.24.542179.

9. Campbell, A., Yim, J., Barzilay, R., Rainforth, T. & Jaakkola, T. Generative flows on discrete state-spaces: Enabling multimodal flows with applications to protein co-design. arXiv [stat.ML] (2024).

10. Varadi, M. et al. AlphaFold Protein Structure Database in 2024: providing structure coverage for over 214 million protein sequences. Nucleic Acids Res. 52, D368–D375 (2024).

11. Cheng, H. et al. ECOD: an evolutionary classification of protein domains. PLoS Comput. Biol. 10, e1003926 (2014).

12. Zhang, J. et al. Predicting protein-protein interactions in the human proteome. Science 390, eadt1630 (2025).

13. Frank, C. et al. Scalable protein design using optimization in a relaxed sequence space. Science 386, 439–445 (2024).

14. Zhang, Y. & Skolnick, J. TM-align: a protein structure alignment algorithm based on the TM-score. Nucleic Acids Res. 33, 2302–2309 (2005).

15. Abramson, J. et al. Accurate structure prediction of biomolecular interactions with AlphaFold 3. Nature 630, 493–500 (05 2024).

16. Kamisetty, H., Ovchinnikov, S. & Baker, D. Assessing the utility of coevolution-based residue-residue contact predictions in a sequence- and structure-rich era. Proc. Natl. Acad. Sci. U. S. A. 110, 15674–15679 (2013).

17. Anishchenko, I., Ovchinnikov, S., Kamisetty, H. & Baker, D. Origins of coevolution between residues distant in protein 3D structures. Proc. Natl. Acad. Sci. U. S. A. 114, 9122–9127 (2017).

18. Fox, D. R., Taveneau, C., Clement, J., Grinter, R. & Knott, G. J. Code to complex: AI-driven de novo binder design. Structure 33, 1631–1642 (2025).

19. Yim, J., et al. Diffusion models in protein structure and docking. Wiley Interdiscip. Rev. Comput. Mol. Sci. 14, e1711 (2024).

20. Goverde, C. A. et al. Computational design of soluble and functional membrane protein analogues. Nature 631, 449–458 (2024).

21. Wicky, B. I. M. et al. Hallucinating symmetric protein assemblies. Science 378, 56–61 (2022).

22. Pillai, A. et al. De novo design of allosterically switchable protein assemblies. Nature 632, 911–920 (2024).

23. Cao, L. et al. Design of protein-binding proteins from the target structure alone. Nature 605, 551–560 (03 2022).

24. Evans, R. et al. Protein complex prediction with AlphaFold-Multimer. bioRxiv 2021–2010 (2021) doi:10.1101/2021.10.04.463034.

25. Introducing BenchBB and the community paper of the Protein Design Competition. https://www.adaptyvbio.com/blog/benchbb (2025).

26. Nooren, I. M. A. & Thornton, J. M. Diversity of protein-protein interactions. EMBO J. 22, 3486–3492 (2003).

27. Janin, J., Bahadur, R. P. & Chakrabarti, P. Protein-protein interaction and quaternary structure. Q. Rev. Biophys. 41, 133–180 (2008).

28. Pace, C. N. & Scholtz, J. M. A helix propensity scale based on experimental studies of peptides and proteins. Biophys. J. 75, 422–427 (1998).

29. Huyghues-Despointes, B. M., Scholtz, J. M. & Baldwin, R. L. Effect of a single aspartate on helix stability at different positions in a neutral alanine-based peptide. Protein Sci. 2, 1604–1611 (1993).

30. Yan, C., Wu, F., Jernigan, R. L., Dobbs, D. & Honavar, V. Characterization of protein-protein interfaces. Protein J. 27, 59–70 (2008).

31. Huising, M. O., Kruiswijk, C. P. & Flik, G. Phylogeny and evolution of class-I helical cytokines. J. Endocrinol. 189, 1–25 (2006).

32. Schwartz, D. M., Bonelli, M., Gadina, M. & O’Shea, J. J. Type I/II cytokines, JAKs, and new strategies for treating autoimmune diseases. Nat. Rev. Rheumatol. 12, 25–36 (2016).

33. Rider, P., Carmi, Y. & Cohen, I. Biologics for targeting inflammatory cytokines, clinical uses, and limitations. Int. J. Cell Biol. 2016, 9259646 (2016).

34. Marks, D. S. et al. Protein 3D structure computed from evolutionary sequence variation. PLoS One 6, e28766 (2011).

35. UniProt Consortium. UniProt: The universal protein knowledgebase in 2023. Nucleic Acids Res. 51, D523–D531 (2023).

36. Sharir-Ivry, A. & Xia, Y. Quantifying evolutionary importance of protein sites: A Tale of two measures. PLoS Genet. 17, e1009476 (2021).

37. Beltrao, P., Bork, P., Krogan, N. J. & van Noort, V. Evolution and functional cross-talk of protein post-translational modifications. Mol. Syst. Biol. 9, 714 (2013).

38. Shehu, A. & Kavraki, L. E. Modeling structures and motions of loops in protein molecules. Entropy (Basel) 14, 252–290 (2012).

39. Jones, S., Marin, A. & Thornton, J. M. Protein domain interfaces: characterization and comparison with oligomeric protein interfaces. Protein Eng. 13, 77–82 (2000).

40. Zambaldi, V., et al. De novo design of high-affinity protein binders with AlphaProteo. Preprint at https://arxiv.org/abs/2409.08022 (2024).

41. Bennett, N. et al. Improving de novo protein binder design with deep learning. Nature Communications 14, 2625–2625 (05 2023).

42. Steinegger, M. & Söding, J. MMseqs2 enables sensitive protein sequence searching for the analysis of massive data sets. Nat. Biotechnol. 35, 1026–1028 (2017).

43. Remmert, M., Biegert, A., Hauser, A. & Söding, J. HHblits: lightning-fast iterative protein sequence searching by HMM-HMM alignment. Nat. Methods 9, 173–175 (2011).

44. Steinegger, M. et al. HH-suite3 for fast remote homology detection and deep protein annotation. BMC Bioinformatics 20, (2019).

45. Kabsch, W. & Sander, C. Dictionary of protein secondary structure: pattern recognition of hydrogen-bonded and geometrical features. Biopolymers 22, 2577–2637 (1983).

46. Pei, J. & Grishin, N. V. AL2CO: calculation of positional conservation in a protein sequence alignment. Bioinformatics 17, 700–712 (2001).

47. Panaretos, V. M. & Zemel, Y. Statistical Aspects of Wasserstein Distances. arXiv [stat.ME] (2018).

48. Sanderson, T., Bileschi, M. L., Belanger, D. & Colwell, L. J. ProteInfer, deep neural networks for protein functional inference. Elife 12, e80942 (2023).

49. Mann, H. B. & Whitney, D. R. On a Test of Whether one of Two Random Variables is Stochastically Larger than the Other. Ann. Math. Stat. 18, 50–60 (1947).

